# Rapid expansion of podoplanin-positive fibroblasts following radiation limits the anti-tumour CD8+ T-cell response to radiotherapy

**DOI:** 10.1101/2025.10.07.680949

**Authors:** Anna Wilkins, Spyridon Makris, Mary Green, Steve Hooper, Anna Mikolajczak, Giovanni Giangreco, Srinivas Allanki, Emma Nye, Emma Connick, Richard Mitter, Sam Cooper, Emily Durie, Malin Pedersen, Amy Burley, Katie O’Fee, Ben O’Leary, Daniel Shewring, Zoe Ramsden, Alan Melcher, Kevin Harrington, Navita Somaiah, Sophie E. Acton, Erik Sahai

**Author notes:** joint senior authors (Sophie E. Acton, Erik Sahai). Corresponding authors contacts: Anna Wilkins, The Institute of Cancer Research, 123 Old Brompton Road, London SW7 3RP, Sophie E. Acton, University College London, Gower St, London WC1E 6BT, Erik Sahai, The Francis Crick Institute, 1 Midland Road, London NW1 1AT.

## Abstract

Radiotherapy is known to cause changes in the tumour stroma which can undermine treatment efficacy. Our understanding of this process has historically centred around effects driven by Transforming Growth Factor-beta (TGF-β) and alpha-smooth muscle actin (α-SMA)+ fibroblasts. Here, we identified a rapid expansion of podoplanin (PDPN)+ fibroblasts following radiotherapy in breast, head and neck and melanoma tumours. This fibrosis was not dependent on TGF-β, but was downstream of a radiotherapy-induced adaptive immune response. CD8+ T-cells entering the tumour after radiation were sequestered at the interface between residual tumour cells and PDPN+ fibroblasts and failed to enter the tumour core. Genetic deletion of PDPN in fibroblasts impacted their cytoskeleton and ability to organise extracellular matrix. This was associated with increased CD8+ T-cell entry and spontaneous tumour regression. Overall, we identify a mechanism whereby PDPN+ fibrosis limits immune-mediated radiation cell kill and demonstrate that disruption of PDPN signalling favours tumour control.

**Significance:** In this study we show that rapid podoplanin (PDPN)+ fibroblast expansion following radiotherapy limits immune-mediated radiation cell kill. Targeting PDPN and associated downstream signalling improves tumour control and is a promising strategy in combination with radiotherapy.

**Graphical abstract:** 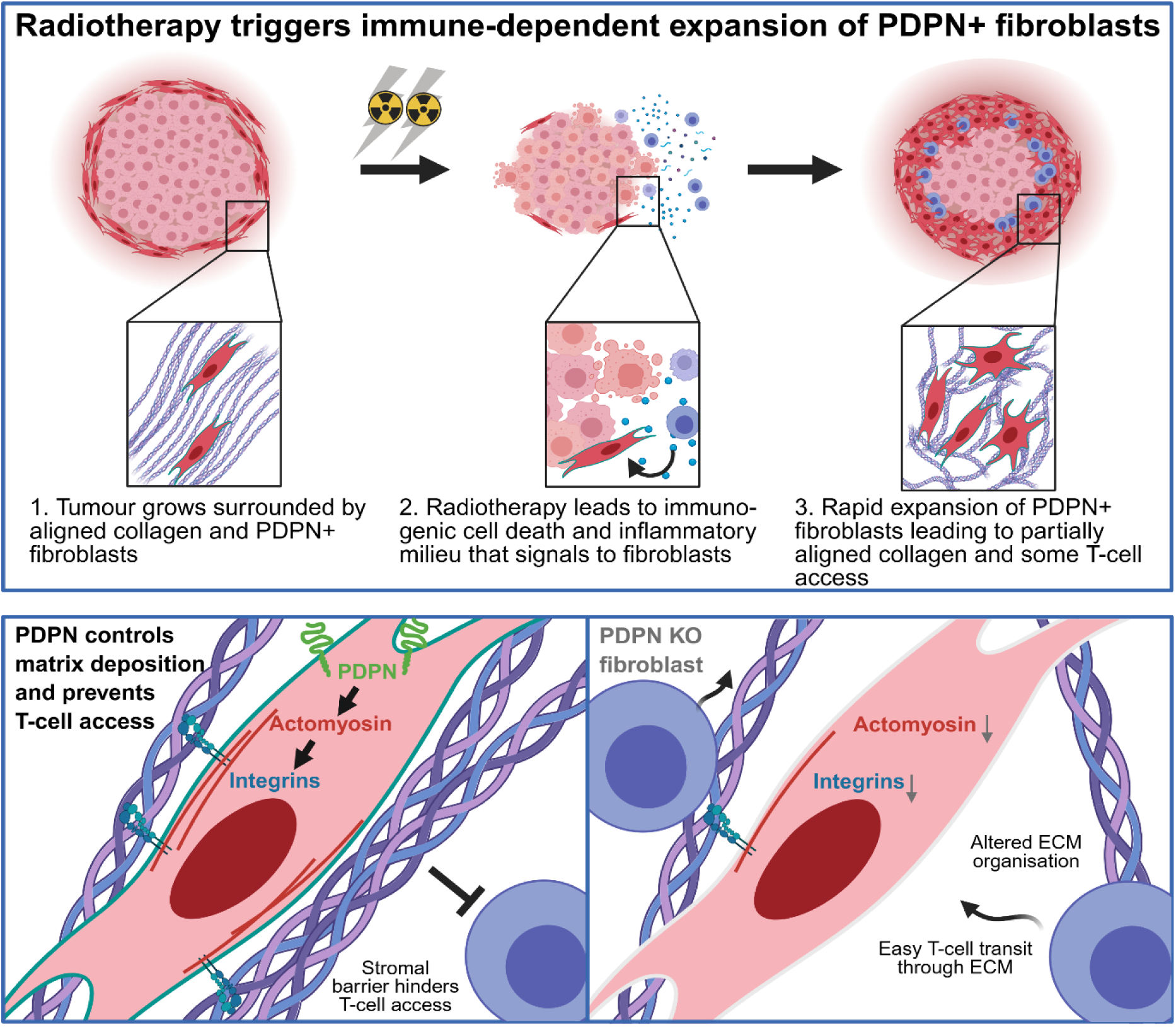

## Introduction

Radiotherapy is a mainstay of cancer treatment, with approximately half of all patients with solid tumours receiving radiotherapy. In addition to the direct induction of lethal DNA damage in cancer cells, radiotherapy also impacts the tumour microenvironment (1). The productive stimulation of the adaptive immune system augments the efficacy of radiotherapy (2). However, the effect of the tumour microenvironment is not always positive. Several studies have implicated the fibroblastic stroma as hindering the response to radiotherapy (3, 4). Transforming Growth Factor beta (TGF-β) promotes the formation of myofibroblast-like cancer-associated fibroblasts, abbreviated as myoCAFs, which typically express alpha-smooth muscle actin (α-SMA) (5). These contractile cells are able to generate a dense extracellular matrix (ECM), which is a common feature of tumours following radiotherapy and linked to unfavourable long-term outcomes (6).

Dense peri-tumoral stroma can lead to the exclusion of T-cells and limit the ability of the immune system to control tumours (7–9). CAFs can also produce various immune-modulating factors, including IL-1β and IL-6 (3, 10–15). These factors can promote the survival of cancer cells and polarise immune cells to a pro-tumorigenic phenotype, which undermine the effectiveness of radiotherapy (16, 17). Thus, it is imperative to understand the interplay between radiotherapy and the tumour microenvironment. In particular, while our understanding of the diversity of CAFs has grown, which CAF subtypes dominate following radiotherapy, and how they might act to modulate the efficacy of the treatment, remains poorly understood. Moreover, apart from TGF-β signalling, there is a paucity of molecular axes that might be targeted to reduce the pro-tumorigenic function of CAFs during radiotherapy.

To address these short-comings, we characterised changes in the tumour microenvironment shortly after radiotherapy in breast and head and neck cancer patients and showed expansion of podoplanin-positive (PDPN+) platelet-derived growth factor receptor-alpha-positive (PDGFRα+) fibroblasts. Next, we established an experimental model of radiotherapy that triggers a large fibroblastic response. Our analyses reveal that the expansion of PDPN+ PDGFRα+ stromal fibroblasts depends on the adaptive immune system, but not TGF-β, indicating that these are not typical myoCAFs. This stromal expansion modulates the access of T-cells into the tumour via two mechanisms: Firstly, ECM deposition during radiation-induced expansion of stromal fibroblasts is not effectively organised into aligned and tightly-packed arrays of fibres, which favours the penetration of CD8+ T-cells. However, this is counteracted by a second mechanism acting via PDPN, which suppresses the recruitment of T-cells and impacts collagen formation in the ECM. Deletion of PDPN in PDFGRα-expressing cells favours tumour elimination.

## Results

### Radiation induces a *PDGFRA+ PDPN+* fibrosis in human breast and head and neck tumours

We evaluated a cohort of 15 patients with breast tumours treated in the KORTUC trial of radiation, who had biopsies before and two weeks after radiotherapy (Figure 1a). We observed widespread fibrotic changes in Haematoxylin and Eosin (H&E) images following radiotherapy (Figure 1b). Following bulk RNA Sequencing of tumour biopsies, we observed that these fibrotic changes were accompanied by significant increases in both *PDPN* and *PDGFRA* genes, but not *ACTA2* (Figure 1c). Widespread PDPN+ αSMA-fibrotic changes were also observed in patients with oropharyngeal head and neck squamous cell cancers who underwent chemoradiotherapy with combination cisplatin and had paired tumour biopsies at baseline and two weeks into radiotherapy (Figure 1d-f and S1A). We also evaluated expression of *PDPN, PDGFRA* and *ACTA2* in published scRNA Seq datasets across ten tumour types including breast cancer and head and neck cancer and confirmed that *PDPN+ PDGFRα+ ACTA2-*fibroblasts were frequently observed (Fig 1g-h and S1b-e) (18–20).

**Figure 1:**
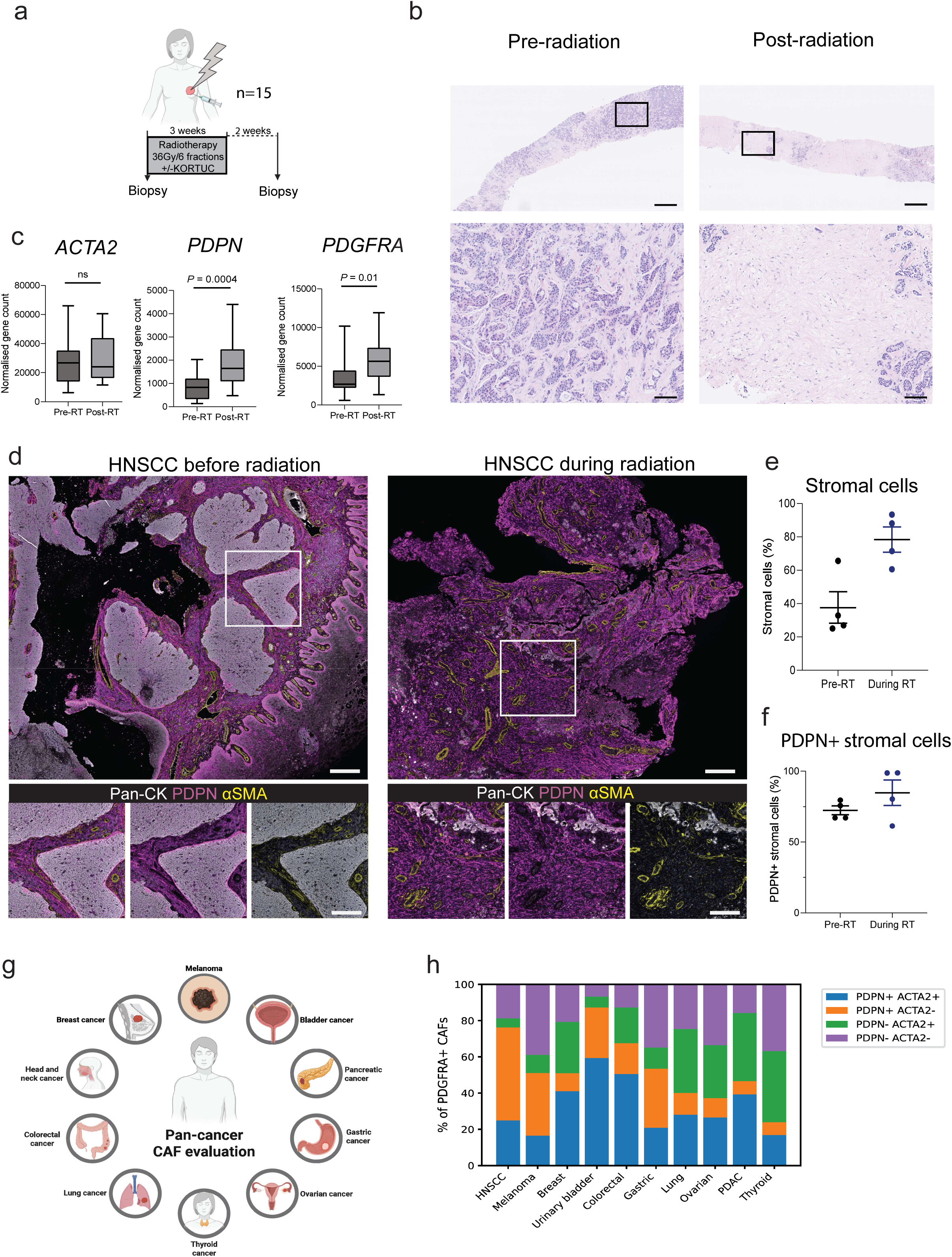
Podoplanin (PDPN+) fibrosis occurs after radiotherapy to breast and head and neck tumours in patients. A: Schematic for pre-and post-radiotherapy biopsies in patient breast tumours. B: Haematoxylin and eosin (H&E) staining to show pre- and post-radiotherapy changes (36 Gy in 6 fractions over 2 weeks, biopsy 2 weeks post radiation). Scale bar top panel 1 mm, bottom panel 100 μm. C: Fibroblast genes *ACTA2, PDPN* and *PDGFRA* before and after radiation (measured using bulk RNA-seq), n=15 paired biopsies. D: Immunofluorescence of head and neck squamous cell carcinoma (HNSCC) before and 2 weeks into radiotherapy (planned treatment 65Gy in 30 fractions over 6 weeks) showing podoplanin (PDPN), alpha smooth muscle actin (α-SMA) and pan-cytokeratin, upper panel scale bar 250 μm, lower panel scale bar 100 μm E: Change in stromal cell density during radiotherapy for HNSCC in paired biopsies, 2 ROI per tumour from 2 patients. F: Change in PDPN+ stroma during radiotherapy for HNSCC in paired biopsies from two patients. G: Schematic for pan-cancer CAF analysis of published scRNA Seq datasets. H: Pan-cancer evaluation of *PDPN* and *ACTA2* expression in *PDGFRA*+ CAFs.

### An expansion of PDPN+ and PDGFRα+ fibroblastic stroma occurs following irradiation in murine models

To understand better these radiation-induced effects on the tumour microenvironment, we established a syngeneic pre-clinical model. *PDPN+ PDGFRα+ ACTA2-*fibroblasts are a prominent feature of the stroma in human melanoma (Fig 1h and SF1b, e) (19), hence we chose a melanoma model whose interplay with the immune system and fibroblastic response to therapy have been extensively characterised – 5555 melanoma cells (21, 22). We observed reduced growth of this model following either single-fraction (8 Gy) or fractionated (3 x 4 Gy) radiotherapy (Fig 2a-b). This was associated with a marked increase in collagen-rich stroma around the tumour one week after radiation (Figures 2c-e). To our surprise, the cells associated with this desmoplastic response were not α-SMA positive, as has previously been observed following radiation (6). Instead, corresponding to our observations in patient tumours, they were positive for the stromal marker, PDPN (Figures 2f-g). We further characterised these cells using multiparameter flow cytometry which enabled us to confirm the PDPN-high, PDGFRα-high and PDGFRβ-low desmoplastic response (Figures 2h-i). Moreover, combining radiotherapy with blockade of TGF-β ligands (using 1D11 monoclonal antibody) had no effect on either the extent of ECM generation or the number or phenotype of CAFs following radiotherapy (Figures S2a and 2h, 2i). Together these data establish that while radiotherapy induces extensive matrix deposition, it is linked neither to the generation of canonical myofibroblasts/myoCAFs nor TGF-β. We observed similar findings of a PDPN+ fibroblast expansion following radiation (2 x 8 Gy) in a second syngeneic pre-clinical model (mEER, head and neck cancer) (Figures S2b-d). Once again, minimal changes in α-SMA were observed (Figures S2c-d).

**Figure 2:**
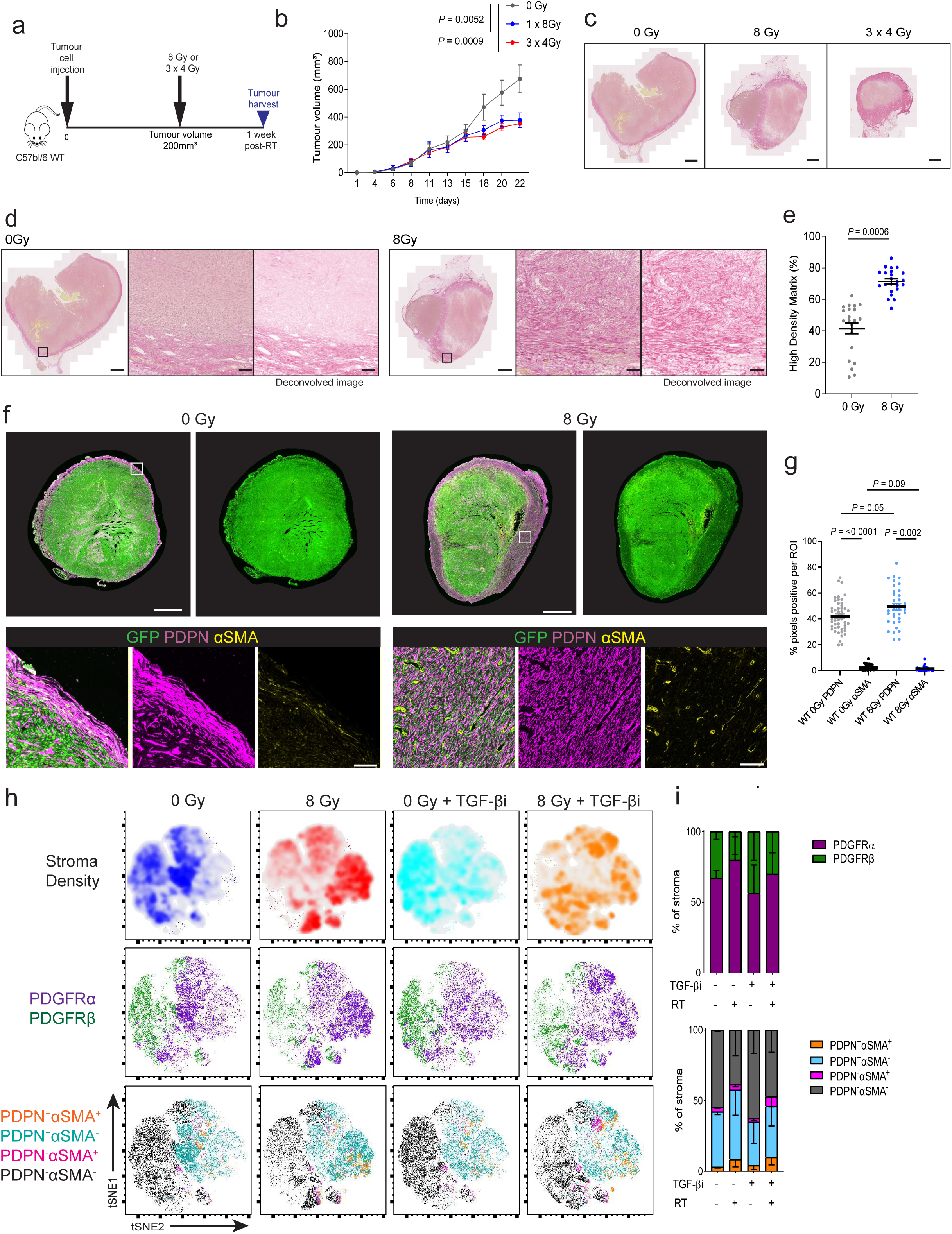
Single-dose and fractionated radiotherapy induces a rapid peri-tumoural fibrotic response. A: Schematic of experimental approach to study of tumour microenvironment post-radiation, B: Growth curves of 5555 tumours treated with 0 Gy, 8 Gy and 3 x 4 Gy, n=10-13 mice per group. C: Picrosirius red (PSR) staining of harvested tumours 1 week following radiation (scale bar 1000μm), D: Methodology for profiling extracellular matrix (ECM) at tumour boundary (scale bar 10μm), including collagen deconvolution, based on PSR staining, E: Quantification of high density matrix using TWOMBLI with and without radiation, 2-3 representative images per tumour, n=7 tumours per group, F: Immunofluorescence of tumours 1 week after 0 Gy versus 8 Gy showing podoplanin (PDPN) and alpha smooth muscle actin (α-SMA), upper panel scale bar 1 mm, lower panel scale bar 100 μm), G: Quantification of PDPN and α-SMA at tumour edge 1 week after 0 Gy or 8 Gy, n=6-8 tumours per group. H: t-distributed stochastic neighbour embedding (tSNE) flow cytometry plots to show changes in stromal fibroblast density with radiation and/or inhibition of Transforming Growth Factor beta inhibitor (TGF-βi) 1D11. I: Quantification of changes in stromal fibroblasts expressing PDGFRα, PDGFRβ, PDPN and α-SMA with radiation and TGF-βi, n=3-5 tumours per group.

### Spatially-distinct inflammatory signalling occurs in PDPN+ fibroblasts following radiotherapy

Long-term control of 5555 tumours following radiotherapy was dependent on the adaptive immune system (Figure S3a and b). In view of this, we next set out to characterise the PDPN+ PDGFRα+ fibroblasts induced by radiotherapy and their interactions with cancer cells and CD8+ T-cells. To this end, we performed NanoString GeoMx analysis concentrating on PDPN-positive, CD8-positive, and GFP-positive regions of 5555 tumours, corresponding to stromal fibroblasts, CD8+ T-cells, and tumour cells, respectively. These analyses were performed on 5555 tumours treated in control, irradiated (8 Gy), anti-TGF-β (1D11-treated), and combination-treated (radiotherapy and anti-TGF-β) groups, with analysis further sub-divided into tumour core, tumour edge, and outer stroma zones (Figure 3a).

**Figure 3:**
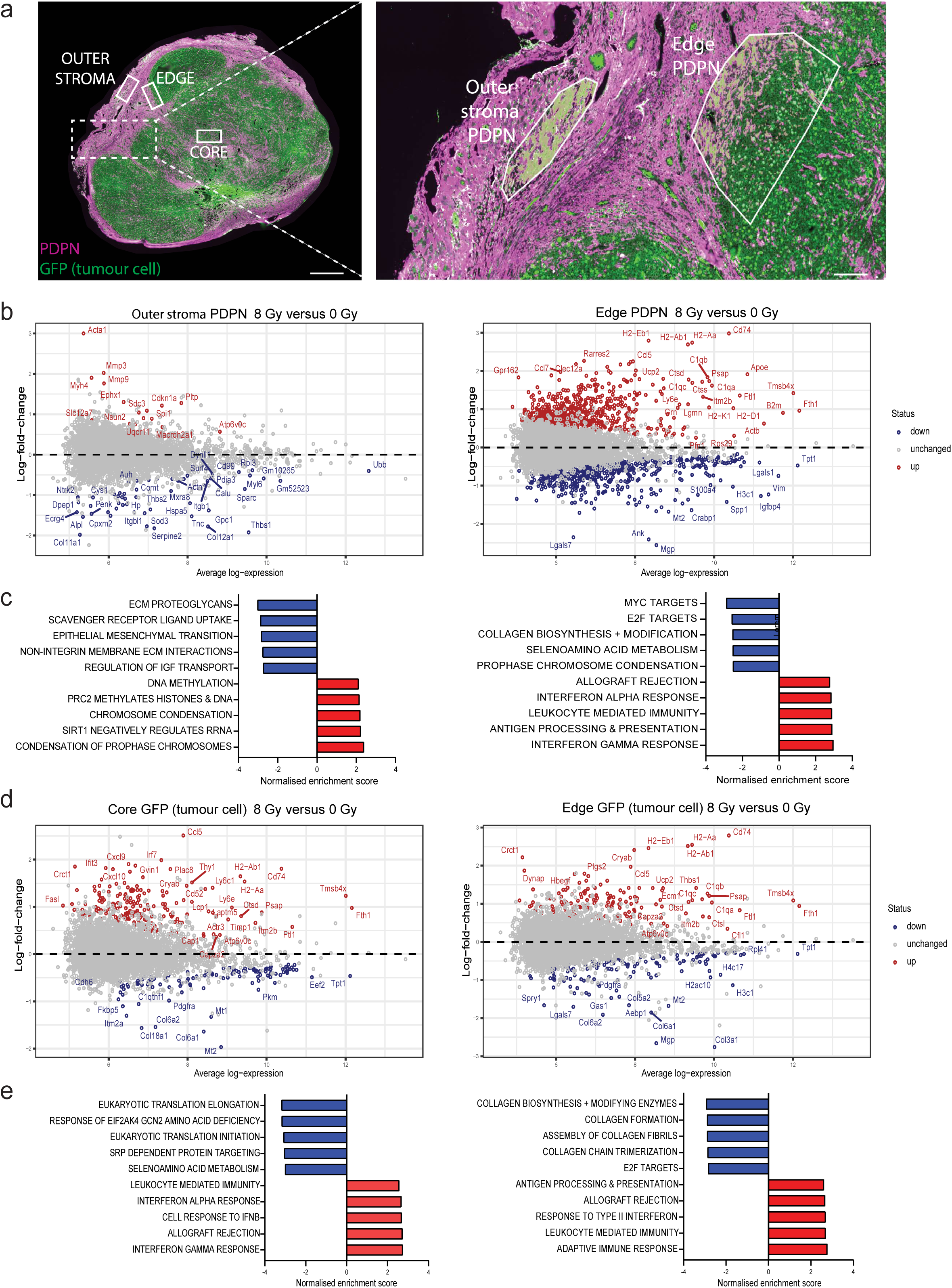
Podoplanin (PDPN)+ stroma shows spatially distinct inflammatory changes with radiation. A: Immunofluorescence of PDPN (fibroblasts) and GFP (tumour cells) on GeoMx platform to show spatially distinct regions of interest (ROI). Left panel scale bar 1 mm, right panel scale bar 200 μm. B: Differential gene expression with 8 Gy (versus 0 Gy) in PDPN+ cells in outer peri-tumoural stroma (left panel) or at residual tumour edge (right panel). 2 tumours, each with 4 ROI, per group C: Pathways showing most significant upregulation or downregulation with 8 Gy (versus 0 Gy) in outer peri-tumoural stroma (left panel) or at residual tumour edge (right panel). D: Differential gene expression with 8 Gy (versus 0 Gy) in GFP+ tumour cells in tumour core (left panel) or at residual tumour edge (right panel) 2 tumours, each with 4 ROI, per group. E: Pathways showing most significant upregulation or downregulation with 8 Gy (versus 0 Gy) in outer peri-tumoural stroma (left panel) or at residual tumour edge (right panel). All genes and pathways shown have significant differences of FDR<0.05.

The peri-tumoral PDPN+ PDGFRα+ stroma induced by radiotherapy showed high levels of expression of interferon-related genes, complement, and antigen-presenting machinery, indicating that these fibroblasts share transcriptional features with previously reported interferon CAFs and antigen-presenting CAFs (Figures 3b-c) (23, 24). These changes were not observed in the outer stroma away from the tumour-stroma boundary (Figures 3b-c and S3c), indicating a possible dependence on interactions with immune or tumour cells, rather than a direct effect of radiation on fibroblasts. Cdkn1a was elevated in fibroblasts away from the boundary, and in the peri-tumoural region (Fig 3b), which likely reflects their response to DNA damage and possible entry into a senescent-like state (25, 26). Consistent with the analysis in Figure 2, the transcriptional profile of the PDPN+ stroma did not show a reduction in typical TGF-β-driven genes following treatment with 1D11 (Figure S3d-e). TGF-β blockade with 1D11 reduced the expression of TGF-β target genes in T-cells, confirming the drug-on-target effects of 1D11 and suggesting that the T cells are influenced by TGF-β present in the tumour microenvironment (Figure S3f-g).

GFP+ tumour cells at the tumour edge and tumour core showed similar transcriptional changes with radiation at both locations (Figure 3d-e). These included significant increases in type I and II interferon responses, as well as genes that are established T cell chemoattractants, including CCL5, CXCL9 and CXCL10 (Figure 3d-e, S3h) (27–30). In combination, these findings indicate that radiation triggers immunogenic cell death in tumour cells which is associated with spatially-distinct inflammatory signals in PDPN+ fibroblasts directly adjacent to tumour cells, but not in fibroblasts at a distance from tumour cells.

### Expansion of PDPN+ PDGFRA+ stroma depends on the adaptive immune system

To understand potential communication between PDPN+ PDGFRα+ CAFs, CD8+ T-cells, and tumour cells, we performed NicheNet analysis on the GeoMx NanoString output (Figure 4a) (31). These data suggested extensive potential bi-directional communication between PDPN+ PDGFRα+ CAFs and CD8+ T-cells at the tumour margin, including interferon gamma signalling from T cells to CAFs (Figure 4b and Figures S4a-c). In accord with this, we observed that interferon-stimulated genes, including *SIRPA, SAMDH, IRF8* and *SAMHD1,* increased following radiation in patient breast tumours (Figure S4d). These data are consistent with reports that radiation-induced cell killing induces interferon responses (32, 33), and raised the possibility that the immune response to the tumour influenced CAFs. To test this, we repeated the irradiation of tumours in Rag1-knockout (KO) mice lacking an adaptive immune response (Figure 4c-d). There was similar tumour control in both wild-type and Rag1-KO mice 7 days after radiation, indicating that the initial effect of radiotherapy is direct killing of tumour cells. However, the desmoplastic response to irradiation and expansion of PDPN+ PDGFRα+ fibroblasts did not happen in Rag1-KO mice (Figures 4e-j). Thus, the desmoplastic response following irradiation is not a direct effect on the fibroblastic stroma, but a response to increased immune surveillance of the tumour, possibly resulting from immunogenic cancer cell death.

**Figure 4:**
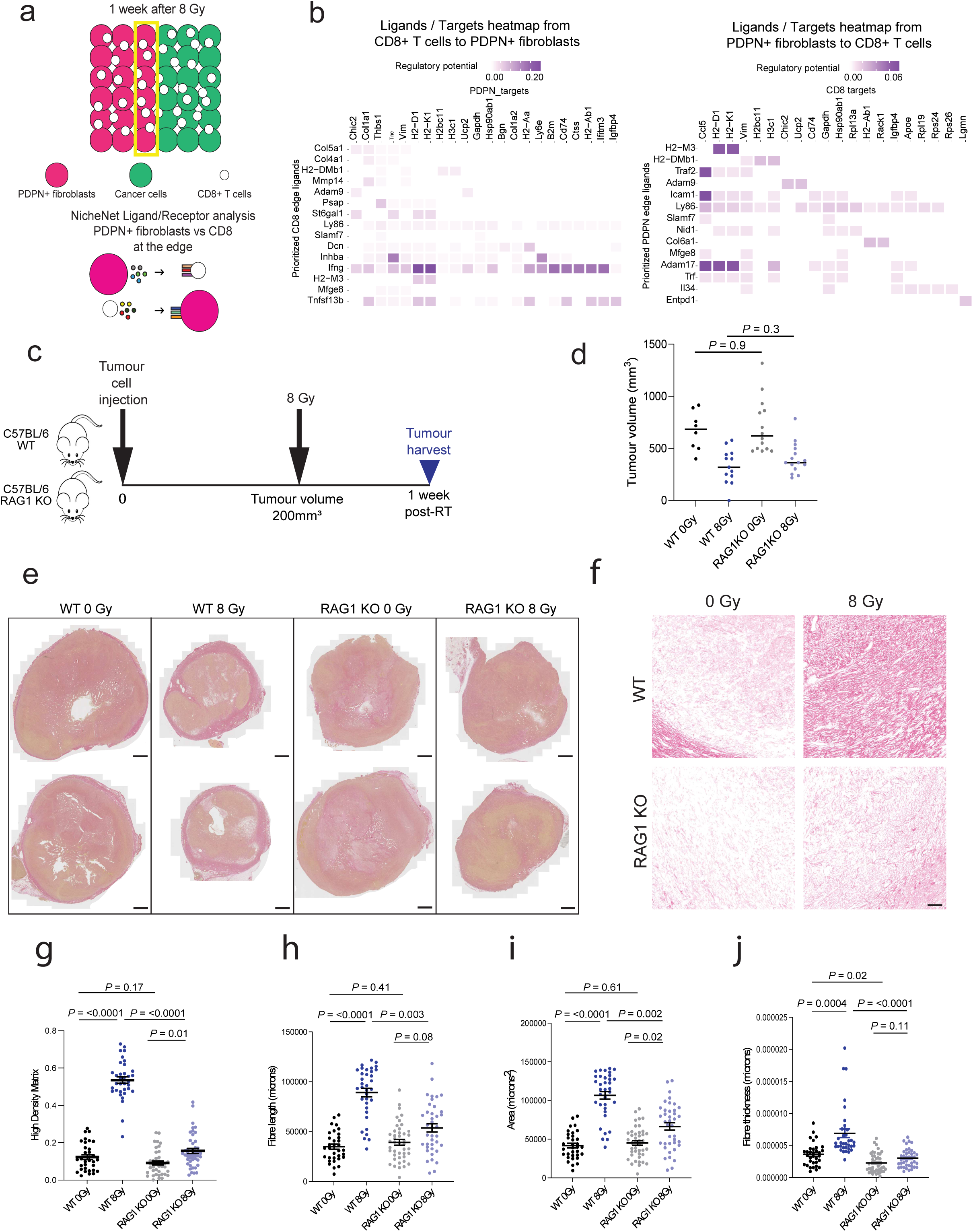
Radiotherapy-induced fibrosis is dependent on T cells. A: Schematic showing NicheNet analysis of ligand receptor interactions between CD8+ T-cells and PDPN+ stroma. B: Ligand/targets heatmap to show top 15 ligands from CD8+ T-cells to PDPN+ stroma cells (left panel) and from PDPN+ stroma cells to CD8+ T-cells (right panel) based on 2 tumours, each with 4 ROI, per cell type. C: Schematic of experimental approach evaluating radiotherapy in wild type and RAG1-KO C57BL/6 mice, D: Tumour size at harvest 1 week following 8 Gy or 0 Gy for tumours grown in WT versus RAG1-KO mice (n=8-15 mice per group), E: Picrosirius Red staining of collagen in WT and RAG1-KO tumours treated with 0 Gy versus 8 Gy (scale bar 1000 μm), F: Exemplar PSR images, with collagen deconvolution, of tumour edge in WT versus RAG1-KO mice treated with 0 Gy versus 8 Gy (scale bar 50 μm), G-J: Quantification of ECM at tumour edge as per images in 1D based on high density matrix, fibre length, area and fibre thickness (n=11-14 tumours per group).

### Radiotherapy alters ECM organisation favouring T-cell access

Given the importance of the adaptive immune system for durable tumour control in our model (Figure S3a,b), and the relevance of CD8+ T-cells implied by the transcriptional analysis, we proceeded to investigate the spatial distribution of T-cells. Following radiation, both multiplex immunofluorescence and flow cytometry demonstrated an influx of CD8+ T-cells, but not CD4+ T-cells (Figures 5a-c). We initially hypothesised that the increased collagen deposition that followed radiotherapy would entirely prevent the entry of T-cells. However, irradiation led to increased accumulation of CD8+ T-cells in the stroma proximal to the tumour boundary, i.e. at the interface between peri-tumoural fibrosis and irradiated tumour cells (Figure 5a). The boundary between collagen and tumour cells was clearly demarcated in all tumours however more detailed examination of the collagen deposited following irradiation revealed that it had collagen fibres that were less aligned than non-irradiated collagen surrounding the tumour (Figure 5d). This observation of loss of alignment following radiation was replicated in *in vitro* assays of fibroblast-derived matrix, where we also observed larger gaps between fibres (Figures 5e-g). This indicates that the change in ECM structure is a direct effect of irradiation on the fibroblasts, not an indirect consequence of signals emanating from other cells in the tumour microenvironment. Thus, there are two opposing effects at play after radiotherapy – the expansion of CAFs creates a thicker stromal barrier, but the more porous nature of the ECM deposited by fibroblasts following irradiation favours access. The net result is accumulation of CD8+ T-cells at the tumour margin.

**Figure 5:**
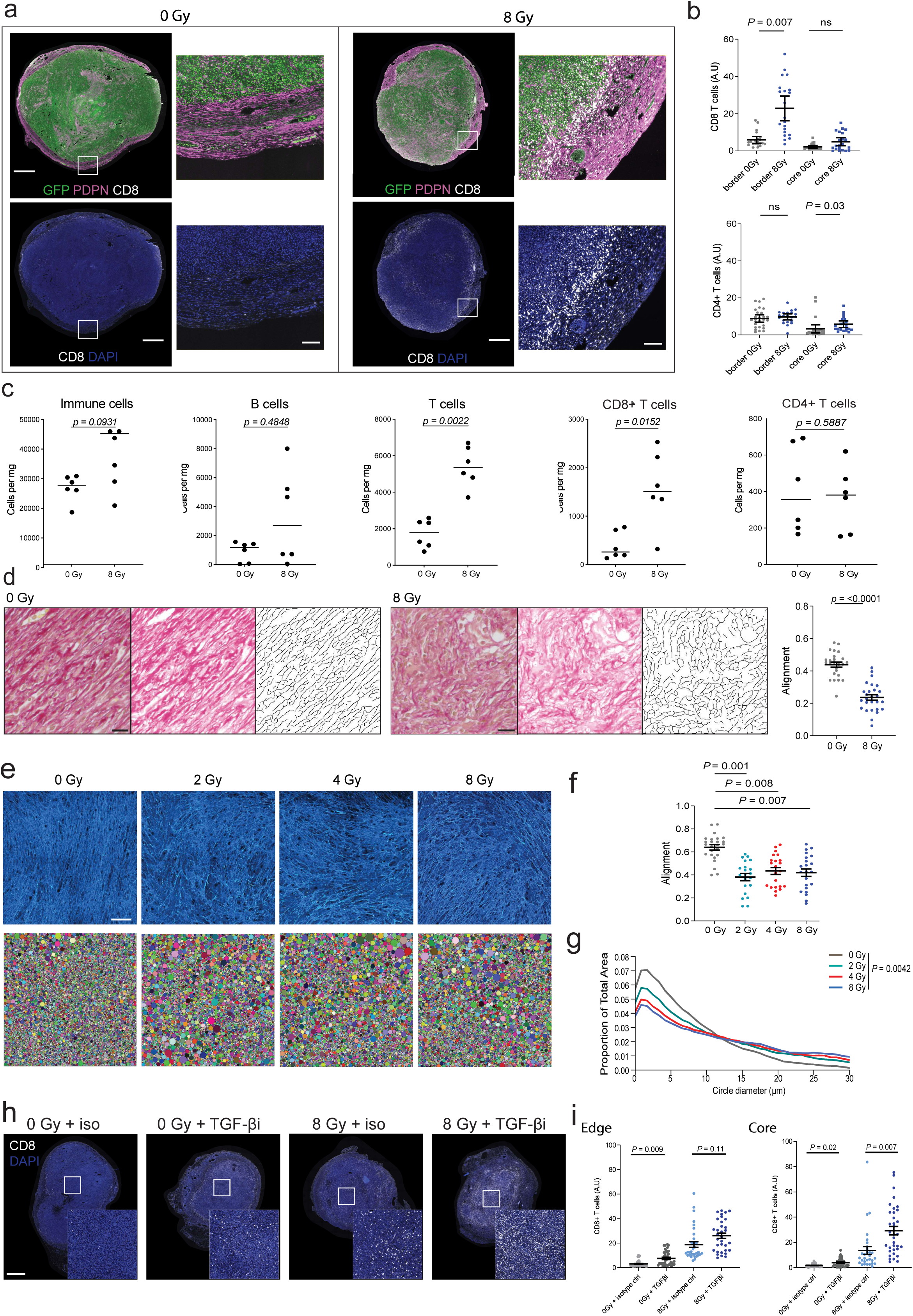
CD8+ T-cells accumulate at tumour/stroma boundary following radiation. A: Immunofluorescence to show CD8+ T-cell localisation at tumour edge (GFP+ tumour cells), scale bar top panel: 1 mm, bottom panel 100 μm B: Quantification of CD8+ T-cell localisation at tumour border and core (top) and CD4+ T-cell localisation (bottom), n=7-9 tumours per group C: Immune cell infiltration evaluated with flow cytometry for Immune cells, B cells, T cells, CD8+ and CD4+ T-cells, n=6 tumours per group. D: Picrosirius red staining of collagen at tumour edge, with and without irradiation, to show change in alignment including quantification, n=6 tumours per group. E: Cell matrix-derived assay to show changes in ECM alignment and pore size. Scale bar 100 μm. F: Quantification of alignment changes in E. G: Quantification of pore size changes in E. H: Changes in CD8+ T-cell distribution with radiation and TGF-βi 1D11. Scale bar 1 mm, I: Quantification of CD8+ T-cells in core and edge of tumours (exemplified in H), n=10-14 tumours per group.

Our transcriptional analysis suggested that T-cells are directly responding to TGF-β (Figure S3g); therefore, we investigated whether this could account for the failure of CD8+ T-cells to move beyond the tumour margin and enter the core of tumours. Figure 5h-i shows that blockade of TGFβ signalling leads to increased entry of CD8+ T-cells into the tumour core, and that this is further enhanced following radiotherapy. Taken together, these data indicate that multiple factors impact the entry of T-cells into tumours following radiation. These include TGFβ signalling acting on CD8+ T-cells to suppress their entry into tumours, and changes in the abundance and organisation of the ECM.

### PDPN functions to reduce T-cell infiltration of the stroma

Having established that irradiation leads to increased T-cell numbers in the stroma surrounding the tumour, but not their effective access to the tumour, we sought to identify the molecular mechanism by which the stroma prevented T-cells from eliminating tumours. PDPN is critical for the ability of fibroblastic reticular cells to modulate immune cell function within lymph nodes, leading us to hypothesise that it may also play an important role in peritumoral fibroblasts (34–38). To test this, we crossed PDGFRα-CreERT2 mice with PDPN-floxed mice thereby generating mice in which PDPN could be deleted specifically from PDGFRα-expressing cells. These mice were then injected with tumour cells before the administration of daily tamoxifen from days 3 to 7 days after injection (Figure 6a). This resulted in tumours that lacked PDPN expression in stromal fibroblasts (Figure 6b and d), and led to an increase in spontaneous rejection of tumours, increasing from 10% to 25% (Figure 6c). Consistent with this, there was a marked slowing of tumour growth between days 11 and 13, which corresponds with the timing of an adaptive immune response to the tumour cells (Figure S6a).

**Figure 6:**
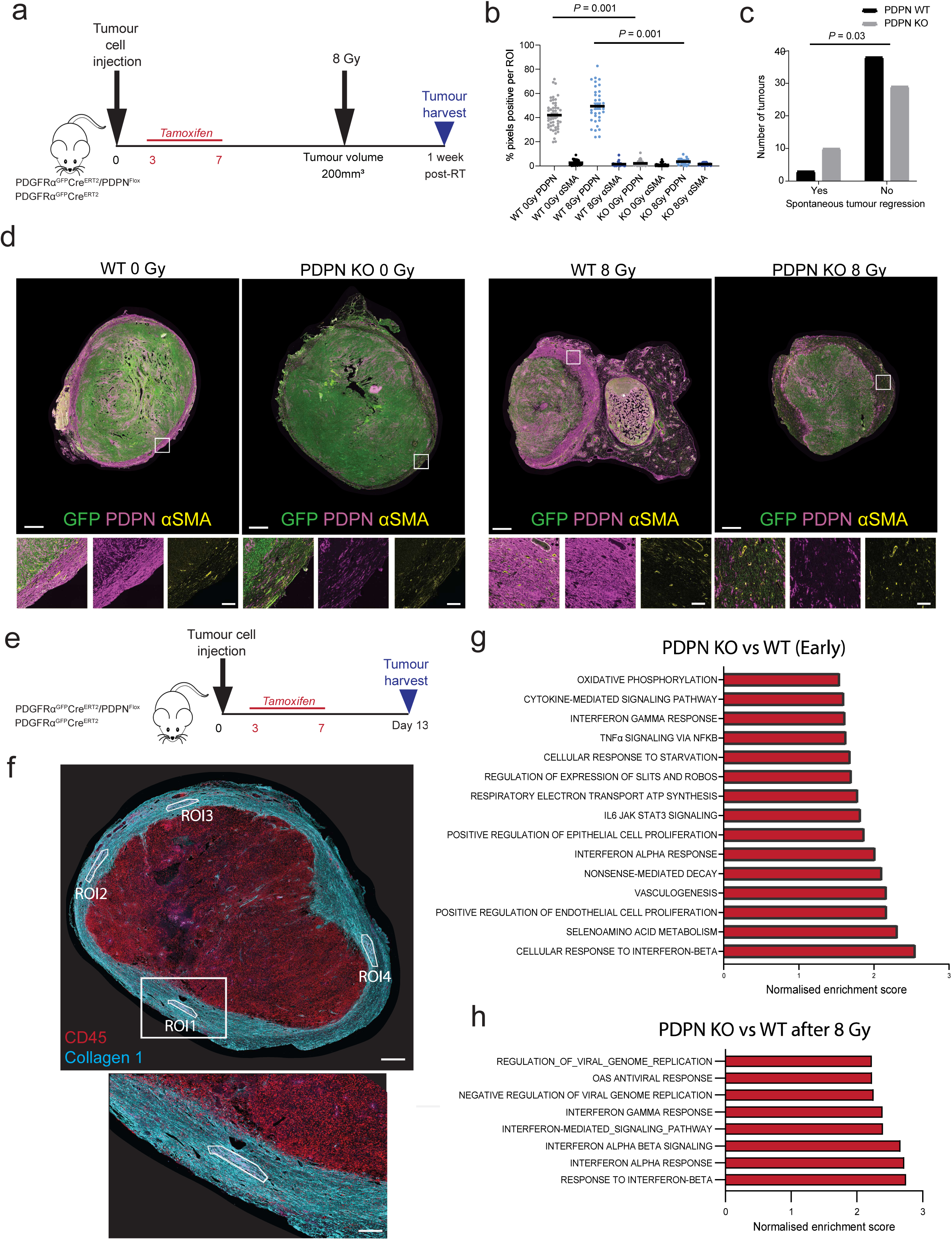
PDPN depletion in PDGFRα+ fibroblasts increases spontaneous tumour regression. A: Schematic of experimental approach evaluating the impact of PDPN-KO in tumours treated with irradiation, B: Quantification of PDPN versus α-SMA+ at tumour edge following KO of PDPN in PDGFRα positive cells (see D), n=4-5 tumours per group KO, n=6-8 tumours per group WT. C: Rate of spontaneous tumour regression (prior to irradiation), n=41 mice WT and 38 mice KO. D: Spatial localisation of PDPN versus α-SMA+ following KO of PDPN in PDGFRα positive cells (scale bar top panel 1 mm, bottom panel 100 μm. E: Schematic of experimental approach evaluating impact of PDPN-KO on early tumour growth (Day 13). F: Immunofluorescence to show ROI selection based on collagen I image masks to identify fibroblasts at tumour boundary. Upper panel scale bar 500 μm. Lower panel 200 μm. G: Pathways upregulated in PDPN-KO versus WT tumours at day 13 post tumour cell injection (4 ROI from each of 4 tumours per group). H: Pathways upregulated in PDPN-KO vs WT tumours 1 week after treatment with 8 Gy (4 ROI from each of 3 tumours per group).

To gain greater insights into the mechanisms underlying the above observations, further tumours were harvested from the above model at day 13 after tumour cell injection (Figure 6e). We then adopted a two-pronged approach in which we used spatial transcriptomics and histological evaluation to profile PDPN-knockout (KO) versus wild-type (WT) tumours. NanoString GeoMx was used to profile transcriptomic signalling of fibroblasts with PDPN-KO versus WT in tumours harvested at both the early and later timepoints described above. The later tumours included those treated with radiation. Collagen I was used to identify fibroblasts in the tumour stroma with additional use of CD45 to subtract signals from neighbouring immune cells (Figure 6f). At early timepoints, PDPN-KO fibroblasts showed increase in type I and II interferon response pathways, as well as other inflammatory pathways including IL6-JAK-STAT3 signalling and TNFα signalling via NFκB (Figure 6g). Of note, a number of these inflammatory pathways have shown similar upregulation in PDPN KO fibroblastic reticular cells (38). Consistent with these observations, interferon response pathways were also upregulated in PDPN-KO fibroblasts one week after 8 Gy of radiation (Figure 6h). These data are consistent with a further increase in CD8+ T-cell signalling to CAFs, (see also interferon signalling from T-cells to CAFs – Figure 4b), and potentially underlie the differences in growth rate and therapy responsiveness shown in Figures S6c-e.

Histological evaluation revealed a significantly increased number of CD8+ T-cells around the tumour edge with PDPN-KO (Figure 7a and b), which is consistent with the elevated interferon signalling in fibroblasts (Figure 6g-h). This is likely to be functionally related to the increased spontaneous tumour regression we observed following PDPN-KO (Figure 6c). The peri-tumoural stroma did not show differences in ECM organisation between WT and KO, however, we identified changes in collagen organisation just inside the tumour cell boundary in tumours with PDPN-KO. Here, the stroma/tumour cell boundary was less clearly defined with longer and more branched fibres penetrating into tumour cell regions, including fibres oriented perpendicular to the tumour boundary (39) (Figures 7c-d and S7a). To explore if these changes reflected a cell-intrinsic change in PDPN-KO fibroblasts, we performed fibroblast-derived matrix assays. These demonstrated that the ability of fibroblasts to organise both collagen and fibronectin into dense networks was directly impacted by PDPN-KO (Figures 7e-f and S7b-c). To explore this phenomenon more deeply, we investigated the cellular machinery involved in fibronectin and collagen fibrillogenesis. Staining for F-actin and active myosin light chain phosphorylated at serine 19 (pS19-MLC), active integrin beta 1 (ITGB1) and fibronectin (FN) revealed that these were markedly disrupted in PDPN KO fibroblasts, with loss of the adhesion points connecting F-actin to fibronectin fibres (Figures 7g-h and S7d-e). To establish if decreased actomyosin function was responsible for the lower levels of integrin activity and fibronectin organisation in PDPN KO cells, we treated wild-type cells with a ROCK inhibitor - Y27632 - that blocks phosphorylation of myosin light chain. Figure S7d-g confirms that ROCK inhibition reduces pS19-MLC; moreover, integrin activity and FN deposition were also reduced following Y27632 treatment. ROCK inhibition did not affect PDPN expression (Figure S7f). These data establish PDPN as an important functional player in stromal fibroblasts that restricts T-cell access to tumours as well as directly impacting integrity of the actomyosin cytoskeleton and ECM architecture.

**Figure 7:**
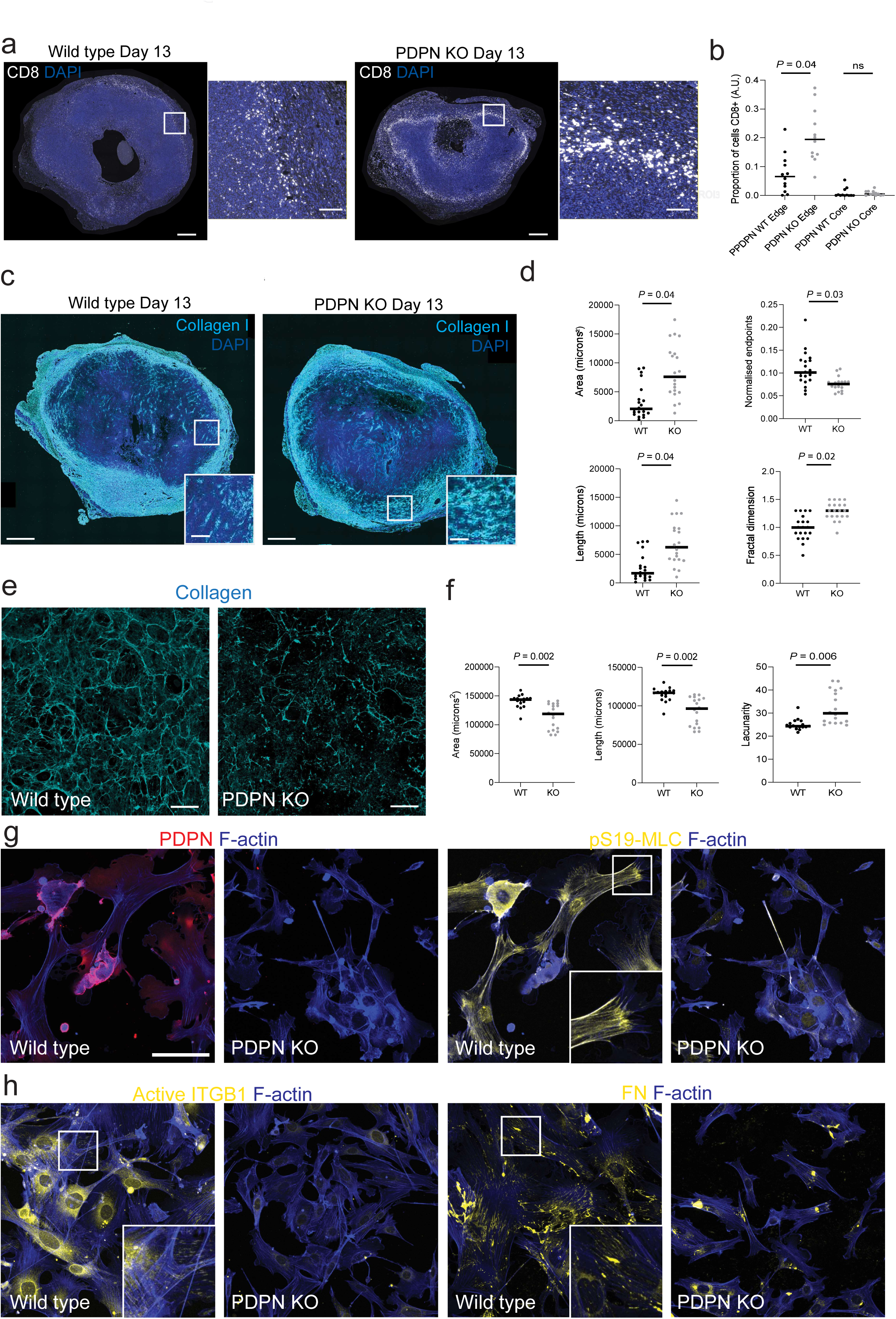
PDPN depletion in PDGFRα+ fibroblasts impacts the extracellular matrix and actin cytoskeleton. A: IF of CD8+ T-cells in PDPN-WT versus -KO. Scale bar large panel: 500 μm, small panel 100 μm. B: Quantification of CD8+ T-cells at tumour edge in PDPN-WT versus KO, n=4 tumour per group. C: IF of collagen in PDPN-WT versus KO at outer tumour cell boundary. scale bar large panel: 500 μm, small panel 100 μm. D: Quantification of collagen patterns inside tumour edge in PDPN-WT versus KO, n=4 tumours per group. E: IF of Collagen in fibroblast-derived matrix (FDM) assays using WT and PDPN-KO murine fibroblasts. Scale bar 100 μm F: Quantification of collagen in FDM in WT versus PDPN-KO contexts. G: IF of fibroblasts from WT versus PDPN-KO mice to show differences in cellular PDPN and intracellular pS19-MLC (myosin light chain phosphorylated at serine 19). Scale bar 50 μm. H: IF of fibroblasts from WT versus PDPN-KO mice to show differences in intracellular active ITGB1 (integrin beta 1) and FN (fibronectin). Scale bar 50 μm.

## Discussion

The identification and targeting of factors limiting the efficacy of radiotherapy has the potential to greatly improve outcomes for patients with cancer. Fibrosis has been known to hamper effective radiotherapy for many years, both by limiting the doses of irradiation that can be delivered safely to normal tissues, and, more recently, by providing a protective environment for residual cancer cells (3, 40, 41). In this work, we established a model with very pronounced generation of a collagen-rich extracellular matrix and the expansion of PDPN+ PDGFRα+ stromal fibroblasts following irradiation. The radiation-induced fibrosis we have identified was observed in breast cancer, melanoma and head and neck cancer. In melanoma, radiotherapy is most commonly used for treatment of *de novo* oligometastatic disease or oligoprogression on systemic therapy; here, high doses per fraction are used in the form of stereotactic radiotherapy. A range of dose schedules are used in the treatment of breast cancer and head and neck cancer. The fibrosis we observed occurred across different fractionation schedules supporting its clinical relevance. Although TGF-β signalling is extensively implicated in fibrosis following radiotherapy (42), we did not observe a role for TGF-β ligands in the response of stromal fibroblasts following radiotherapy. Nonetheless, we did observe changes in CD8+ T-cell distribution and transcriptional state, which suggests TGF-β ligands may signal directly to T-cells in our model, as has been observed elsewhere (29).

The lack of explicit TGF-β signalling to stromal fibroblasts in our model is consistent with the absence of canonical α-SMA high fibroblasts and may suggest that other ligands and cues dominate in instructing changes in fibroblast biology. Of particular interest is the observation of inflammatory signalling between T-cells and fibroblasts following irradiation, and the lack of stromal response in Rag1-KO mice. These data place the immune response to radiotherapy upstream of fibroblast expansion. We propose that fibrosis following irradiation primarily represents a wound healing response to immunogenic tissue damage. In the context of incomplete tumour control, this response further restricts the anti-tumour response by sequestering effector immune cells within the fibrotic tumour boundary outside of tumour cell nests. Specifically, we propose the following model of changes after tumour irradiation. First, immunogenic cell death triggered by damage to cancer cells leads to an inflammatory immune response (Graphical Abstract – step 1). The resulting cytokine milieu, which we infer to be rich in interferons, promotes the expansion of PDPN+ PDGFRα+ fibroblasts that share transcriptional similarity with interferon CAFs and antigen-presenting CAFs (Graphical Abstract – step 2). These fibroblasts deposit extensive ECM, but it is less densely packed than in non-irradiated tumours and permits the influx of T-cells to the tumour periphery (Graphical Abstract – step 3). However, even following irradiation, CAFs and their associated ECM form a clearly demarcated boundary around the residual tumour that hampers CD8+ T-cell access to the tumour core. Thus, the net effect of the stromal expansion resulting from engagement of the adaptive immune system is to hinder elimination of the tumours. In addition, TGF-β signalling in CD8+ T-cells also hampers their ability to enter the tumour core. We propose that this acts as an additional mechanism to limit CD8+ T-cell entry into the tumour core.

Interestingly, PDPN is not just a marker of stromal fibroblasts following irradiation, but it has a critical functional role. Fibroblasts lacking PDPN exhibit defects in collagen organisation in simple cell culture systems (Graphical Abstract – lower panels), which is consistent with previous reports in fibroblastic reticular cells (43). Moreover, we observe defects in the cytoskeletal machinery required for fibronectin fibril generation, termed the fibripositor, that is in turn important for collagen fibre generation (44). We additionally establish that PDPN-dependent regulation of actomyosin activity is a key mechanism controlling integrin activity, which is critical for fibripositor function, and fibronectin assembly. Targeting this response, potentially via pathways downstream of PDPN signalling in fibroblasts, has the potential to improve the therapeutic index of radiotherapy.

Our findings indicate that fibrotic reactions post-radiation can be driven by different CAF subtypes with varying temporal dynamics. We observed an early fibrosis that occurred within one week; this differs markedly from “classical” radiation-induced fibrosis where scar tissue that is thought to be biologically indolent forms over a number of months. This early fibrosis can shape how tumour cells are sensed and killed by the immune system after radiation and hence is closely related to overall radiation efficacy. We propose that a more nuanced consideration of early, middle and late fibrotic effects after radiation is required. A better understanding of early fibrotic events that impact therapy provides the potential for novel therapeutic intervention. Such interventions may have associated impacts on normal tissue fibrosis, and a key consideration is favourably shifting the therapeutic index in terms of both efficacy and radiation-induced side effects.

## Methods

### Patient Samples

Breast tumour samples were obtained from the randomised phase 2 KORTUC trial with ethical approval granted by the West of Scotland Research Ethics Committee (REC reference 20/WS/0019) with full informed consent from the patients. Breast tumour samples were obtained prior to treatment and 2 weeks after last radiotherapy treatment. Radiation was given at a dose of 6 x 6 Gy in a standard linear accelerator at 6-10MV) with or without intra-tumoural hydrogen peroxide as per the KORTUC protocol (45). All patients and investigators were blinded to randomised group. For RNA extraction 3-10 x 10μm sections (depending on tumour content) were obtained from the formalin-fixed paraffin-embedded (FFPE) breast tumour core biopsies. Extraction was performed using the Qiagen RNeasy FFPE kit as per protocol with an additional double ethanol wash and a double elution in the final step. Libraries were constructed with Agilent SureSelect XT HS2 RNA Library, SureSelect XT HS Human All Exon V8 probes were used to capture the entire exome region >35Mb, the library was sequenced using the Illumina Novaseq. For analysis, Bcl2fastq software was used for converting raw basecalls to fastq files and FASTQC was used to assess sequencing data quality. Analysis was performed using STAR alignment software to align to the reference genome (GRCh38), HTSeq-count was used to count the number of mapped reads. In DESeq2 normalised counts were obtained using DESeq2 normalisation method (counts(dds, normalised=TRUE)). Wilcoxon tests (corrected by Benjamini Hochberg for number of genes analysed) were used to compare normalised counts of pre-radiation and post-radiation groups. H&E sections were scanned using a Nanozoomer© (Hamamatsu) digital slide scanner at 40X magnification. Head and neck cancer samples were obtained from the CHIMERA study with ethical approval granted by Camden & Kings Cross Research Ethics Committee (18/LO/1250) with full informed consent from the patients. Baseline archival tissue was obtained with further research biopsies of oropharyngeal tumours taken two weeks into chemo-radiation (65 Gy in 30 fractions with concomitant cisplatin). Immunofluorescence on HNSCC tumours was performed as outlined below.

### scRNA Seq re-analysis

Publicly available data were downloaded from Gene Expression Omnibus (Melanoma - GSE215121, HNSCC - GSE164690, and Pan-cancer CAF data - GSE210347). For the Melanoma dataset, patient samples corresponding to acral Melanoma and cutaneous Melanoma were reanalysed. For the HNSCC dataset, samples corresponding to CD45 negative cells from all the patients available were reanalysed. For the Pan-cancer CAF dataset, the already reanalysed Seurat object was downloaded and the samples were separately reanalysed per tumour type. In general, for all three datasets, each tumour type was analysed as follows: All the patient samples corresponding to each tumour type were analysed together as per the standard methodologies using Scanpy (version 1.10.2). Potential doublets as predicted by the Scrublet package were filtered out. Sample integration was performed using the *scanpy.external.pp.harmony_integrate* function with the patient IDs as the *batch key*. Cells expressing less than 200 genes and greater than 8000 genes were filtered out to remove low quality cells and possible doublets. Cells with mitochondrial content greater than 20% were also filtered out. Principal component analysis (PCA) was performed and the top 20 PCs were used to generate the neighbourhood graph, followed by UMAP embedding. The cancer-associated fibroblast (CAF) clusters were identified using the lineage marker *PDGFRA* expression. CAFs were then subclustered to assess the proportions of the cells expressing *PDGFRA*, *PDPN*, and *ACTA2* (expression value>0).

### Cells and Probes

BRAF-mutant mouse melanoma (5555) cell lines which incorporated an ERK FRET biosensor (EKAREV-NLS) that enabled GFP antibodies to unambiguously detect melanoma cells were established as described previously (22, 46). Melanoma cells were cultured at 37°C in Dulbecco’s modified Eagle’s medium (DMEM) (Invitrogen), supplemented with L-glutamine, 10% foetal bovine serum/1% PenStrep (GIBCO). The murine HNSCC cell line mEER, was cultured similarly and supplemented with 10% FBS (Gibco), 2.4mM L-glutamine, 60mg/mL penicillin, 100mg/mL streptomycin and 0.1mg/mL Primocin (Invivogen #ant-pm-1). Human melanoma-associated fibroblast 1 (MAF1) fibroblasts were isolated from a patient metastatic melanoma and immortalised with lentiviral hTERT as described in Gaggioli, C., et al (47). These patient samples were collected at the Royal Marsden Hospital under ethical approval 15/EE/0151. Cells were selected using 400 μg/mL hygromycin and maintained in DMEM, 10% FCS (PAA Labs) and 1% ITS (insulin–transferrin– selenium; #41400-045; Invitrogen) supplement. PDPN WT and KO CAFs were obtained from *Pdgfrα*-mGFP-CreERT2 x Pdpn^fl/fl^ (Pdgfrα^mGFPΔPDPN^) mouse models (see below). They were immortalised with lentiviral E6. Cells were selected using puromycin and maintained as per MAF1 above. All cells were screened for mycoplasma before culture.

### In vivo experiments

Animal experiments were performed in accordance with UK regulations under project license PPL/80/2368 and approved for the Francis Crick Institute and Institute of Cancer Research by the institutional animal ethics committee review board. For melanoma experiments, 0.2 × 10^6^ 5555 cells were suspended in 200 μL (100μL PBS and 100μL growth factor reduced Matrigel (Sigma-Aldrich) and subcutaneously injected in the right flank of C57BL/6 mice (either wild type, genetically modified with RAG1-KO or conditional deletion of PDPN in cells expressing PDGFRα+ (see below)). For mEER head and neck tumour experiments, 1 × 10^6^ cells were suspended in 100 μL PBS and injected subcutaneously in the right flank of 7-10 week-old female C57Bl/6 wild type mice purchased from Charles River Laboratories.

*Pdgfrα*-mGFP-CreERT2 x Pdpn^fl/fl^ (Pdgfrα^mGFPΔPDPN^) mouse models were generated as described by Horsnell et al (36). Activation of the Cre recombinase was achieved through the administration of tamoxifen (20 mg/mL) resuspended in ethanol/sunflower oil (1:10 ratio). Tamoxifen was dosed at 200 µg/g to a total maximum of 4 mg by oral gavage on 5 consecutive days 3 days following tumour cell injection. Females and males aged 8 - 15 weeks were used for experiments except the mEER model when 7-10 week-old female wild type C57 Bl/6 mice were used. Animals were assigned experimental groups at random when tumours reached threshold size for irradiation (melanoma 200 mm^3^ and 50-55 mm^3^ for head and neck tumours).

1D11 or isotype control (2BScientific BE0057 and BE0083) were diluted in the recommended dilution buffer (2BScientific IP0070 and IP0065) and administered via intra-peritoneal injection at a dose of 10 mg/kg 3 times per week starting 24 hours prior to irradiation/control for 8 days, until up to 24 hours before tumour harvesting.

### Murine irradiation and tumour measurement

#### Melanoma

Radiation was applied when tumours reached a volume of approximately 200 mm^3^. Mice were anaesthetised with intraperitoneal injections of Fentanyl (0.05 mg/kg), Midazolam (5 mg/kg) and Medetomidine (0.5 mg/kg), then treated with either single dose (1 x 8 Gy) or fractionated radiation (3 x 4 Gy delivered on consecutive days) (200 kV, 10 mA, 1 mm Cu filtration), targeted to the flank. Lead shielding (3 mm thickness) with a 1.5 cm diameter circular aperture was used to target radiation dose to the flank. The machine was calibrated daily to achieve a consistent total dose. The dose rate was approximately 1.4 – 1.5 Gy/min for each treatment, with a radiation time of approximately 5.5 minutes for 8 Gy and 3 minutes for 4 Gy. Probes confirmed radiation outside the aperture was negligible prior to experiments. Anaesthetic reversal was carried out using intraperitoneal injections of Naloxone (1.2 mg/kg), Flumazenil (0.5 mg/kg) and Atipam (2.5 mg/kg). Recovery was performed in warming chambers at 37°C. Mouse body weights and tumours were measured three times a week using callipers and tumour volume calculated by V = (W^2^ × L)/2.

#### Head and neck

Radiation was applied when tumours reached a volume of approximately 50 mm^3^. The animals were irradiated under anaesthesia with Xylazine / Ketamine given by intraperitoneal injection. Local tumour radiotherapy was delivered in two fractions of 8 Gy each on consecutive days (day 13-14) using a AGO250 kV X-ray machine using lead shielding. Radiation dose was measured using a Farmer Chamber and Unidos-E Dosimeter (PTW). Diet gel was provided after radiotherapy sessions. Mouse body weights and tumours were measured two times a week and tumours volumes were calculated using the formula length x width x height (mm) x 0.5236.

### Fibroblast-derived matrix (FDM) assay and in vitro fibroblast immunofluorescence

The FDM assay was performed as described in Franco-Barraza et al (48). In brief, 24-well glass-bottom dishes (MatTek, P35-1.5-14-C) were pre-prepared with 0.2% gelatin solution for 1 hour at 37 °C, followed by 1% glutaraldehyde for 30 minutes at room temperature. The plate was washed twice with PBS buffer solution, then incubated with 1M ethanolamine for 30 minutes at room temperature. The plate was washed twice with PBS before seeding 8×10^4^ cells in medium supplemented with 100 μg/ml ascorbic acid ((+)-sodium L-ascorbate, Sigma, A4034). The cells were maintained for 5 days and the medium changed every 2 days. Cells were removed using the extraction buffer described and washed several times with PBS before undertaking immunofluorescence for fibronectin using anti-fibronectin (1:1,000 dilution; Sigma, F3648) and anti-fibronectin-FITC (1:50 dilution; Abcam, ab72686). Collagen was profiled using the CNA35 genetically-encoded collagen binding probe labelled with Tdtomato (49) (dilution 1:500). Immunofluorescence for P-MLC was performed using rabbit 3671L Cell Signalling Technologies (dilution 1:500). IF for active ITGB1 was performed using rat 9EG7 BD Biosciences (dilution 1:500).

### Imaging and quantification of extracellular matrix

Picrosirius red staining (Abcam picrosirius red stain kit) was used to visualise collagen in FFPE tumours except for PDPN KO versus WT tumours where collagen I was used as part of GeoMx Digital Spatial Profiling. Extracellular matrix patterns in murine tumours, including high density matrix, were quantified using the FIJI plug in TWOMBLI (50). For FDM, CTFire was used to quantify alignment (51). Following image pre-processing, including despeckling and particle filling, pore size was quantified using the Max Inscribed Circles FIJI plug in with a threshold of 2 pixels or 1μm.

### Immunofluorescence/immunohistochemistry imaging and analysis

All multiplex staining protocols were completed using the Leica BOND Rx automated immunostainer. This was programmed to complete the following steps: 1. tissue dewaxing and rehydration, 2. antigen retrieval, 3. primary antibody staining, 4. secondary antibody staining, 5. application of Opal and all appropriate washes. Steps 2-5 were repeated for a further 4 cycles, with an additional protease treatment before Collagen I using RNAscope™ 2.5 LS Protease III (ACD Bio-techne). Slides were counterstained with DAPI and coverslipped before imaging on the PhenoImager HT (Akoya) at 20X followed by spectral unmixing with inForm software (Akoya) and restitching in QuPath v0.4.1 for further analysis. Mouse and human antibody panels for multiplex staining are shown in supplementary methods tables S1 and S2 respectively.

Analysis of immunohistochemistry and immunofluorescence was performed in QuPath v0.4.1. Using the pixel classifier tool, the software was trained based on defined PDPN+ and α-SMA+ regions. A pixel classification was then created and applied to all peri-tumoural fibrotic regions. CD4+ T cell and CD8+ T-cell quantification was performed from immunohistochemistry images where the software was trained to detect positive versus negatively stained cells, the algorithm was then applied to the tumour/stroma interface and tumour core.

### NanoString GeoMx Digital Spatial Profiling (DSP)

Radiation + TGF-βi experiment: NanoString GeoMx Digital Spatial Profiler platform, was used to profile 12 different FFPE tumours (+/- 8 Gy with 1D11 TGF-βi or isotype control, 3 tumours per group). Slides were prepared according to manufacturer’s instructions including heat retrieval, protease treatment (0.01% ProK), probe incubation, immunostaining with GFP, CD8 and PDPN with fluorescently conjugated secondary antibodies and Syto13 nuclear counterstain before imaging on the GeoMx. For each tumour, 4 ROIs were selected in each of 3 different spatial locations, tumour core, tumour edge and distant stroma per tumour where tumour size permitted this. Within each ROI, segmentation masks were defined by visual threshold for T-cells (CD8+), tumour cells (GFP+) and stromal cells (PDPN+) using antibodies shown in supplementary table S3.

PDPN-KO versus -WT DSP experiment: 4 ROI were selected at the tumour edge of 16 FFPE tumours that were PDPN-KO or -WT (8 early timepoint and 12 late timepoint). Collagen I was used to mask stromal cells with CD45 staining negatively selected to exclude immune cells from analysis (see Table S3).

Following library preparation, sequencing was performed using the Illumina NovaSeq 6000.

### Bioinformatics analysis of DSP outputs

GeoMx datasets were exported from the NanoString DSP Analysis Suite using the GeoMx NGS Pipeline tool (version-2.3.3.10) for separate analysis using the Bioconductor packages standR [1] and GeoMxWorkflows [2]. Experiment 1 slide 2 was discarded due to consistently low alignment rates (<50%). Experiment 2 segments with alignment rates <70% were discarded. Genes detected in fewer than 10% of segments based on a limit of quantification (LOQ) defined by negative control probes were discarded. Segment count data were normalised using the Trimmed Mean of M-values (TMM) method. Unwanted technical variation due to a slide-effect was estimated using the RUV4 approach (k=1) (52), calculated from 300 negative control genes with the smallest coefficient of variation. Lowly expressed genes were removed using the filterByExpr function. Differential expression analysis between tumour groups was performed using a linear modeling approach via the voomLmFit function from the EdgeR package (53). The model included the slide weight estimate as a covariate and accounted for repeated sampling from the same tumor using a blocking factor. Statistical significance was assessed using eBayes.

Fast Gene Set Enrichment Analysis (FGSEA) (54) was used to assess whether gene sets of known function (MSigDB Hallmarks, MSigDB C2 Reactome and GOBP) were significantly associated with differential expression using a permutation test (n=2000). Genes were pre-ranked based on their logFC in each pairwise comparison. Nichenet analysis was performed on NanoString DSP output.

### Flow cytometry

Tumours were weighed and placed into RPMI 1640 medium on ice and processed as previously described (34–36). Samples were placed into a digestion buffer containing collagenase D (250 mg mL^-1^) (Millipore Sigma, cat# 11088866001), dispase II (800 mg mL^-1^) (Thermo Fisher Scientific, cat# 17105041) and DNaseI (250 mg ml^-1^) (Sigma-Aldrich, cat# 10104159001). Tissue was digested at 37°C and cell suspension was collected every 10 minutes until fully digested. Cells were centrifuged at 350 *g* for 10 minutes and resuspended in PBS containing 1% BSA, mM EDTA, 0.05% NaN_3_ (FACS Buffer). Cells were then filtered in 70 mm filters and counted. A single cell suspension of 2.5 x 10^6^ cells was incubated for 20 minutes at 4°C with a purified rat IgG2b anti mouse CD16/CD32 receptor antibody (Fc Block) (Biolegend, Cat# 10302, RRID: AB_466390). Lymphocytes were surface stained with fluorochrome-conjugated antibodies as shown in supplementary table S4. For surface stain of non-immune cells fluorochrome-conjugated antibodies were used as shown in supplementary table S5.

Cells were surface stained for 25 minutes at 4°C in FACS Buffer. Cells were incubated with fixable viability kit – Zombie Aqua (BioLegend, cat# 423102, RRID: AB_2869122) in PBS for 30 minutes at 4°C. For intracellular staining, cells were fixed and permeabilised using the FOXP3 Fix/Perm Buffer (Biolegend, cat#421403) used according to manufacturer’s protocol. Immune cells were stained with Ki67-PE/Cyanine7 (BD Biosciences, cat# 561283, RRID: AB_10716060) and non-immune cells with Ki-67-BUV737 (BD Biosciences, cat# 567130, RRID: AB_2916461), ACTA2-rabbit anti-mouse (Proteintech, cat# 14395-1-P, RRID: AB_2223009), CD54/ICAM-1-BUV395 (BD Biosciences, cat# 740222, RRID: AB_2739970), CD106/VCAM-1-BUV615 (BD Biosciences, cat# 752534, RRID: AB_2917524) or CCL21-AF700 (R&D Systems, cat#IC457N, RRID: AB_3655881) for 30 minutes at room temperature in permeabilization buffer. Non-immune cells were blocked with Fc Block in permeabilization for 20 minutes at room temperature and then incubated with F(ab’)_2_ Goat anti-Rabbit IgG (H+L), QdotTM605 (Invitrogen, cat# Q11402MP, RRID: AB_10375594) in permeabilization buffer for 30 minutes at room temperature. Samples were analysed on BD LSRFortessa x-50 (FACS Symphony) equipped with 100-nW 405-nm, 100-mW 488-nm, 150-mW 561-nm, 100-mW 637-nm and 60-mW 355-nn lasers with an ND1.0 filter in front of the FSC photodiode. Data was collected on the FACSDIVA software (version 7). Flow cytometry data was analysed using FlowJO Software (Tree Star).

### Statistical analysis

All in vitro experiments were conducted with at least 3 independent biological replicates. Numbers of mice and tumours per experiment are shown in the figure legends. Comparisons between two groups were conducted using either Mann Whitney U tests for non-parametric data or students t-test for parametric data. Comparisons between multiple groups were conducted using ordinary one-way ANOVA. Tumour growth curves were compared using two-way ANOVA or mixed effects models unless stated otherwise. Statistical comparison of sequencing data is described above.

## Data availability

Spatial transcriptomics data are available at GSE304902. Publicly available scRNA Seq data are available from Gene Expression Omnibus (Melanoma - GSE215121, HNSCC - GSE164690, and Pan-cancer CAF data - GSE210347). Other data is available by request to the corresponding authors.

## Funding and Acknowledgements

This manuscript represents independent research supported by the National Institute for Health Research (NIHR) Biomedical Research Centre at The Royal Marsden NHS Foundation Trust and the Institute of Cancer Research, London. The views expressed are those of the author(s) and not necessarily those of the NIHR or the Department of Health and Social Care. The authors acknowledge funding from Cancer Research UK RadNet at The Institute of Cancer Research and The Royal Marsden Hospital and City of London RadNet.

A.W. is funded by a CRUK Clinician Scientist Fellowship (RCCCSF-Nov24/100003).

S.E.A. is funded by Cancer Research UK Careers Development Fellowship CRUK-A19763, Rosetrees Trust Foundation PGS22 100023-CD1 and Cancer Research UK Senior Fellowship CRUK - RCCSCF- May22\100001.

E.S. is supported by the Francis Crick Institute, which receives its core funding from Cancer Research UK (CC2040), the UK Medical Research Council (CC2040), and the Wellcome Trust (CC2040) and the European Research Council (ERC Advanced Grant CAN_ORGANISE, Grant agreement number 101019366).

## Conflicts of interest

A.W. reports research funding from Artera AI, AstraZeneca, Roche Genentech, Veracyte and speaker honoraria from Johnson and Johnson. E.S. reports grants from Novartis, Merck Sharp Dohme, AstraZeneca and personal fees from Phenomic outside the submitted work. GG receives funding from Merck Sharp & Dohme Corp, New Jersey, USA (LKR190557). AB receives PhD funding from AstraZeneca. BOL reports research funding from Pfizer, consultancy and/or speaker honoraria from Pfizer, Merck Serono, Merck, Eisai, Oliver Wyman, Outrun and funding for travel/meeting attendance from Merck Serono and Pfizer.

