## Supplementary Data for "Rapid expansion of podoplanin-positive fibroblasts following radiation limits the anti-tumour CD8+ T-cell response to radiotherapy"

Supplementary Figure 1

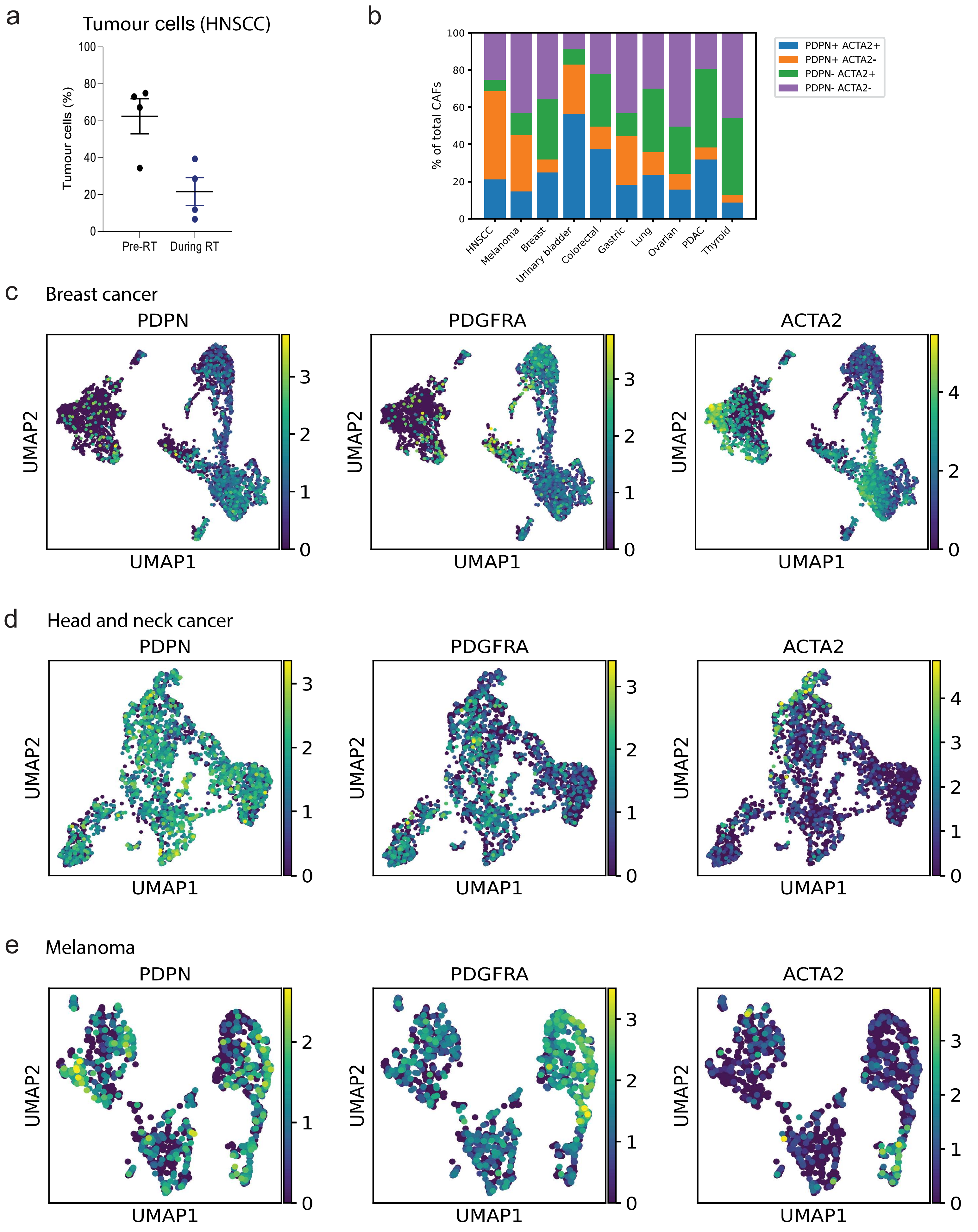

**Supplementary Figure legends**

Supp Figure 1 A: Change in tumour cell density during radiotherapy for HNSCC in paired biopsies, 4 ROI from 2 patients. B: Pan-cancer evaluation of *PDPN* and *ACTA2* expression in all CAFs C-E: UMAP plots to show expression of *PDPN, PDGFRA* and *ACTA2* in fibroblasts from human breast tumours (C), human head and neck cancer (D) and human melanoma (E).

Supplementary Figure 2

a

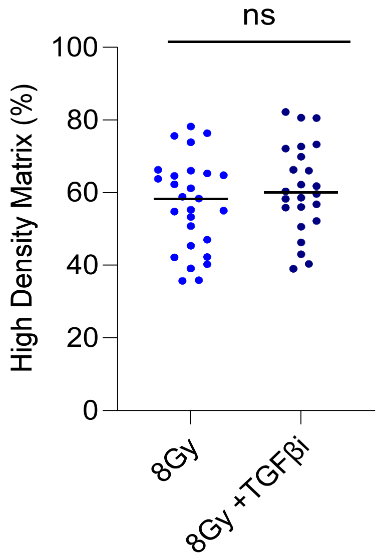


b

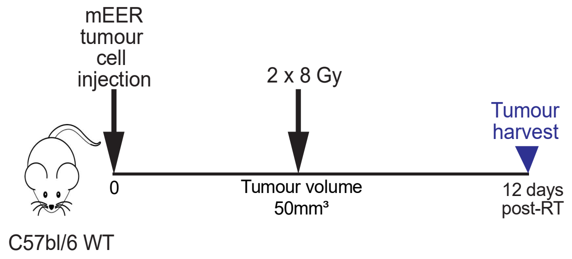


c

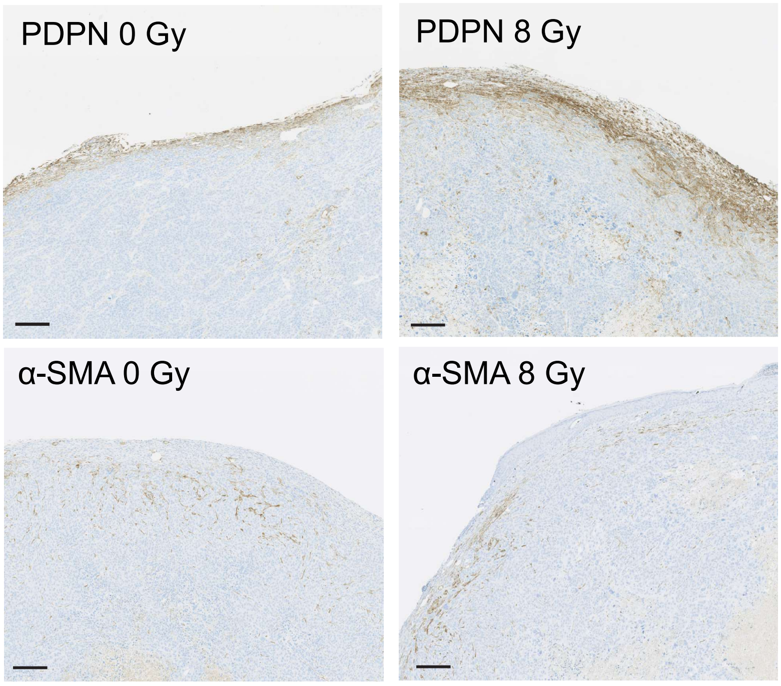


d

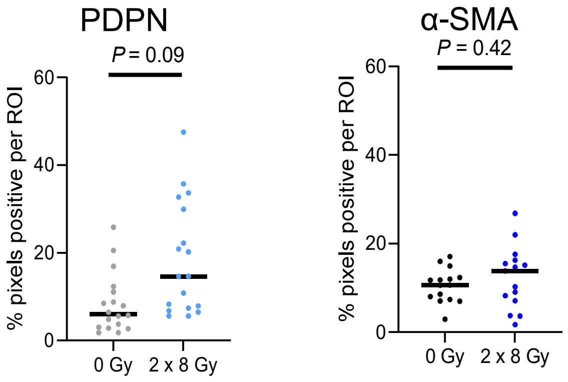

Supp Figure 2 A: Quantification of high density matrix in collagen with and without radiation or inhibition of TGF-β using TWOMBLI. B: Schematic of experimental approach to study effects of radiation to mEER tumours in WT C57 BL/6 mice. C: Immunohistochemistry of PDPN and α-SMA in mEER tumours 14 days after treatment with 2 x 8 Gy or 0 Gy. Scale bar 200 μm. D: Quantification of PDPN and α-SMA at tumour edge in tumours treated with 2 x 8 Gy versus 0 Gy, PDPN n=6 tumours per group, α-SMA n=5 tumours per group.

Supplementary Figure 3

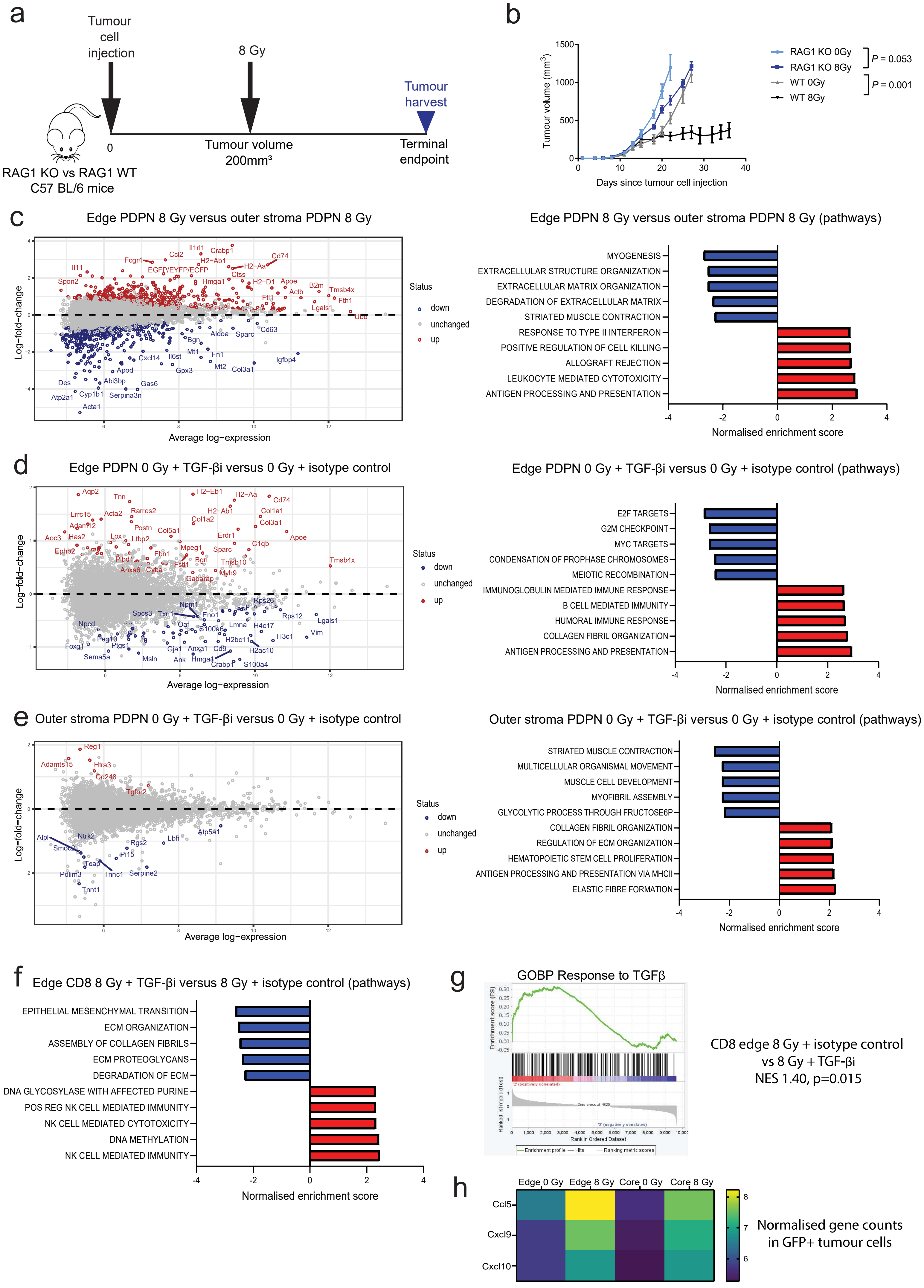

Supplementary Figure 3 A: Schematic of experimental approach to study effects of radiation of RAG1-KO versus -WT C57 BL/6 mice, B: Growth curves of 5555 tumours treated with 0 Gy or 8 Gy in RAG1-KO versus -WT C57 BL/6 mice, n=7-12 mice per group, statistical comparisons are performed at day 22 for RAG1-KO and day 27 WT. C: Differential gene expression with 8 Gy in PDPN+ cells at residual tumour edge versus in outer peri-tumoural stroma and pathways showing most significant upregulation or downregulation with 8 Gy in PDPN+ cells at residual tumour edge versus in outer peri-tumoural stroma, 2 tumours, each with 4 ROI, per group. D: Differential gene expression with TGF-βi in PDPN+ cells at residual tumour edge versus control and corresponding pathways showing most significant upregulation or downregulation. 2 tumours, each with 4 ROI, per group. E: Differential gene expression with TGF-βi in PDPN+ cells in outer peri-tumoural stroma versus control and corresponding pathways showing most significant upregulation or downregulation. 2 tumours, each with 4 ROI, per group. F: Differential gene expression with TGF-βi plus 8 Gy in CD8+ T-cells versus isotype control plus 8 Gy at tumour edge (left hand panel) versus tumour core (right hand panel), 2 tumours, each with 4 ROI, per group. G: Pathway analysis for GOBP for Response to TGF-β comparing CD8+ T-cells at edge treated with 8Gy plus isotype control versus 8 Gy plus TGF-βi Normalised enrichment score (NES) 1.40, p=0.015. All genes and pathways shown have significant differences of FDR<0.05. H: Heatmap showing changes in genes *CCL5, CXCL9* and *CXCL10* in GFP+ tumour cells with radiation according to location.

### Supplementary Figure 4

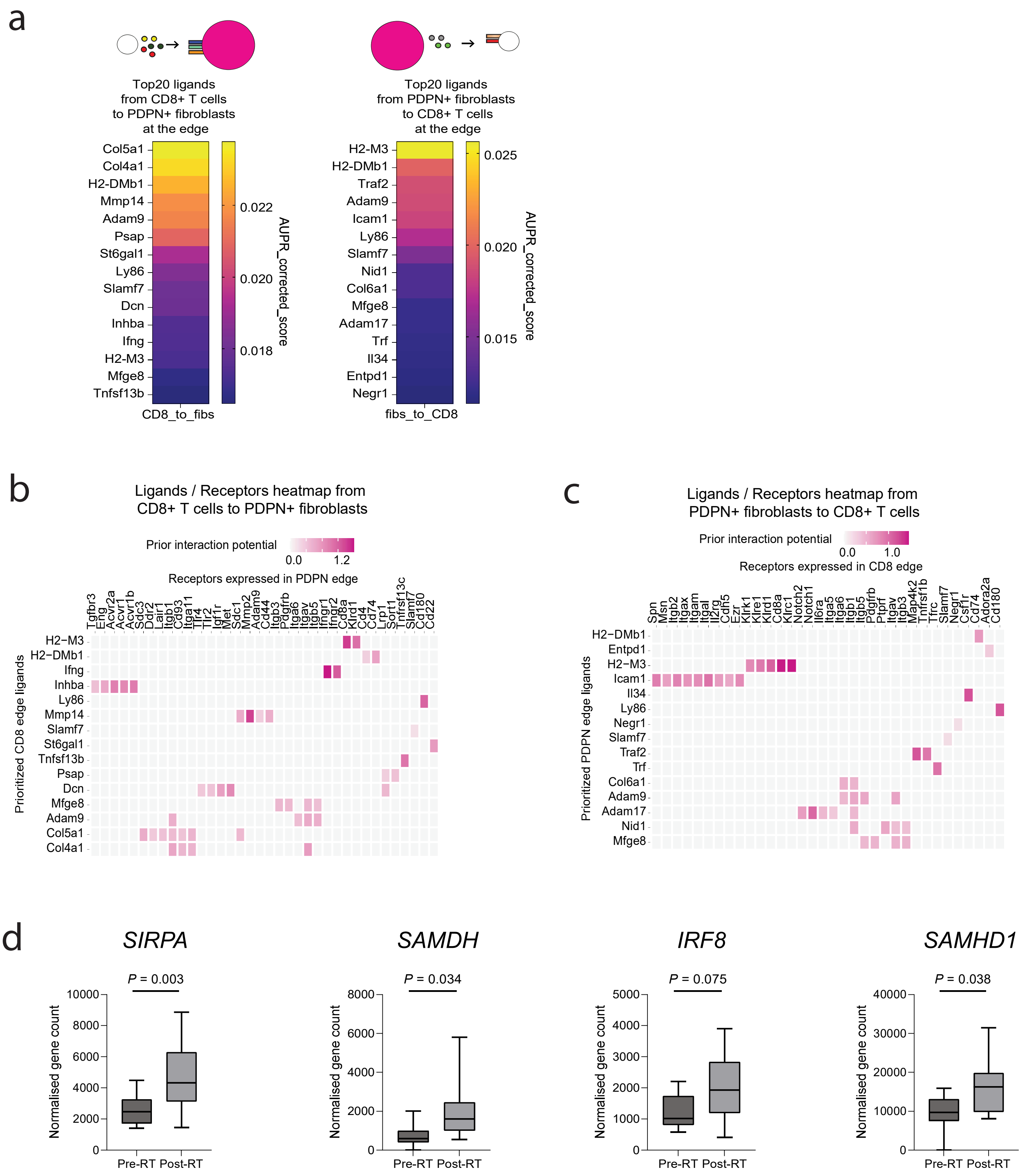

Supplementary Figure 4 A: Heatmap showing top 15 ligands from CD8+ T-cells to PDPN+ stroma cells (left panel) and from PDPN+ stroma cells to CD8+ T-cells (right panel). B: Ligand receptors heatmap to show top 15 ligands from CD8+ T-cells to PDPN+ stroma cells (left panel) and C: from PDPN+ stroma cells to CD8+ T-cells (right panel). D: Interferon-stimulated genes *SIRPA, SAMDH, IRF8* and *SAMHD1* before and after radiation to patient breast tumours (measured using bulk RNA-seq).

### Supplementary Figure 5

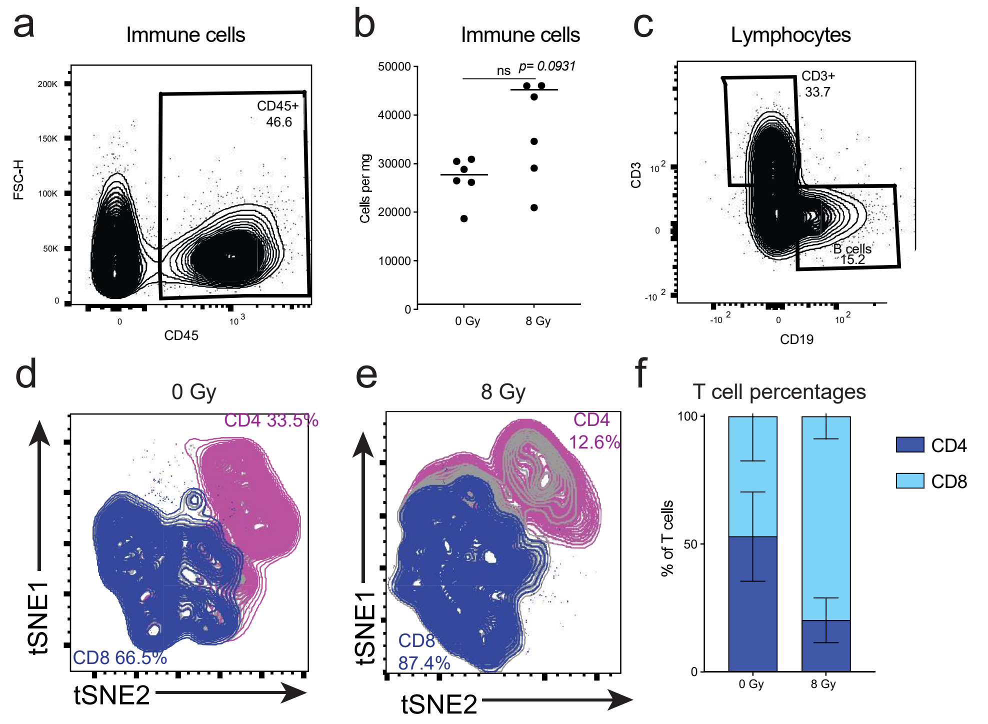

Supplementary Figure 5 A-C: Immune cell infiltration evaluated with flow cytometry for Immune (CD45+) cells (flow density map A and quantification B) and lymphocytes (CD3+ or CD19+) (C), n=6 tumours per group. D-F: T-cell infiltration according to CD8 and CD4 staining. Flow density map 0 Gy (D) and 8 Gy (E). F: Quantification of D and E, n=6 tumours per group.

### Supplementary Figure 6

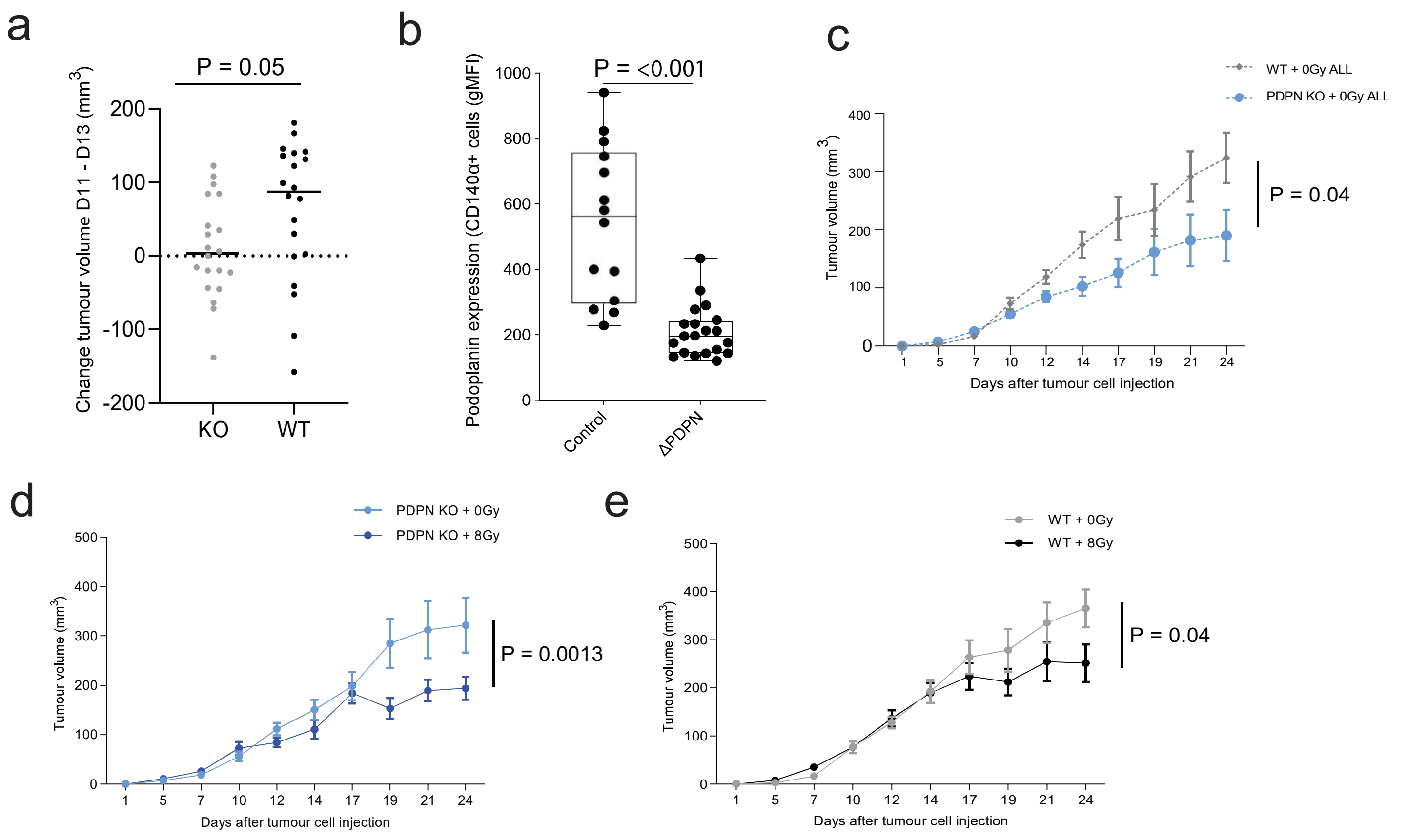

Supplementary Figure 6 A: Change in tumour volume between days 11 and 13 following PDPN-KO or in WT 5555 tumours, n=20 mice per group. B: Flow cytometry to shown PDPN expression in fibroblasts at day 13 after tumour cell injection, n=17-20 mice per group. C: Tumour growth kinetics in all untreated PDPN-WT vs -KO tumours including tumours which did not reach size for randomisation, n=23-26 mice per group. D: Tumour growth kinetics in PDPN-KO tumours treated with 0 Gy versus 8 Gy, n=12-15 mice per group E: Tumour growth kinetics in PDPN-WT tumours treated with 0 Gy versus 8 Gy, n=18-20 mice per group.

Supplementary Figure 7

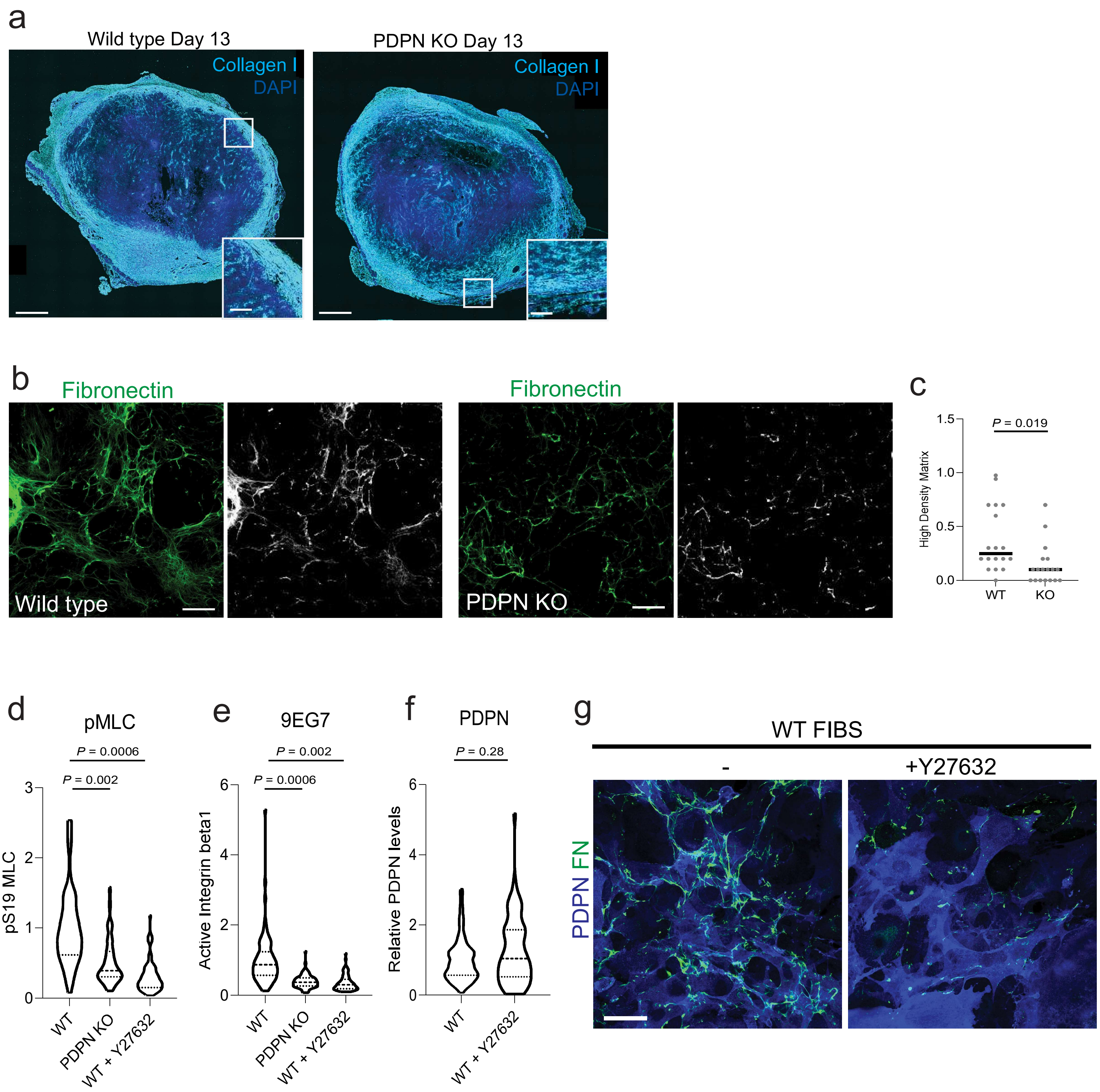

Supplementary Figure 7 A: IF of collagen in PDPN-WT versus KO showing tumour outer edge. scale bar large panel: 500 µm, small panel 100 µm. B: IF of fibronectin in FDM using WT and PDPN-KO murine fibroblasts with corresponding high density matrix image masks. C: Quantification of fibronectin in FDM in wild type versus PDPN-KO contexts. Scale bar 100 µm. D and E: Quantification of active myosin light chain phosphorylated at serine 19 (pS19-MLC) and active integrin beta 1 (ITGB1) in PDPN WT and PDPN KO fibroblasts, including WT fibroblasts with Y27632 ROCK inhibitor. F: Quantification of PDPN in WT fibroblasts with and without Y27632 ROCK inhibitor. G: Immunofluorescence to show impact of Y27632 ROCK inhibitor on fibronectin and podoplanin in WT fibroblasts. Scale bar 100 µm.

#### Supplementary Methods

Table S1: Multiplex immunofluorescence mouse panel

| Antibody sequence | Antibody | Catalogue number | Primary Dilution | Pre-treatment | Opal TSA Reagent | Opal Dilution |
|---|---|---|---|---|---|---|
| 1 | CD8 | 14-0808-82 | 1:200 | ER1 95˚C 20min | 620 | 1:200 |
| 2 | GFP | A-6455 | 1:1000 | ER1 95˚C 20min | 480 | 1:500 |
| 3 | α-SMA | M0851 | 1:250 | ER1 95˚C 20min | 520 | 1:500 |
| 4 | Podoplanin | ABIN 115147 | 1:2000 | ER1 95˚C 20min | 570 | 1:500 |
| 5 | Collagen I | AB758 | 1:200 | ER2 95˚C 20min + ACD protease III 40˚C 15min | 690 | 1:200 |

Table S2: Multiplex immunofluorescence human panel (HNSCC)

| Target | Catalogue number | Antibody dilution | Abundance | Pre-treatment | Opal | Opal brightness | Final position |
|---|---|---|---|---|---|---|---|
| PDPN | Biolegend 916605 D2-40 | 1:800 | High | pH 6.0 95˚C 20min | 620 | Low | 4 |
| αSMA | Abcam ab240654 1A4 | 1:5000 | High | pH 6.0 95˚C 20min | 520 | Medium | 5 |
| panCK | Dako M3515 AE1/AE3 | 1:200 | High | pH 6.0 95˚C 20min | 780 | Lowest | 6 |

Note FAP, CD8 and PDGFRα occupied positions 1, 2 and 3 respectively, data not shown.

Table S3: Antibodies used in GeoMx spatial transcriptomics

| Marker | Catalogue number | Dilution | Secondary | Catalogue number |
|---|---|---|---|---|
| Syto13 | S7575 | 1:50 | N/A | N/A |
| GFP | 2955 | 1:1000 | donkey anti-mouse IgG AF555 | A32773 |
| CD8 | ab217344 | 1:500 | donkey anti-Rabbit IgG AF594 | A21207 |
| Podoplanin | ABIN115147 | 1:2000 | goat anti-syrian hamster IgG AF647 | A21451 |
| CD45 | 70275S | 1:100 | donkey anti-Rabbit IgG AF555 | A-31572 |
| Collagen I | AB758 | 1:100 | Donkey anti-Goat IgG AF647 | A-21447 |

Table S4: Fluorochrome-conjugated antibodies used in surface staining of lymphocytes for flow cytometry

| | |
|---|---|
| CD45-BV750 | BioLegend, cat# 103157, RRID: AB2734155 |
| CD19-PE | BioLegend, cat# 115508, RRID: AB_313643 |
| CD3-AF700 | BD Biosciences, cat# 561388, RRID: AB_10642588 |
| CD4-PE-CF594 | BD Biosciences, cat# 562285, RRID: AB_11154410 |
| CD8a-BUV805 | BD Biosciences, cat# 564920, RRID: AB_2716856 |
| CD44-BV605 | BD Biosciences, cat# 563058, RRID: AB_2737979 |
| CD62L-BUV395 | BD Biosciences, cat# 569400, RRID: AB_3685037 |
| CTLA4-PerCP/Cyanine5.5 | BioLegend, cat# 106316, RRID: AB_2564474 |
| TCF1 | BD Biosciences, cat#566692, RRID: AB_2869822 |
| IL-7Ra-BV711 | BioLegend, cat# 135035, RRID: AB_2564577 |
| CD25-APC | BioLegend, cat# 102012, RRID: AB_312861 |
| PD-1-APC/Fire$^{TM}$810 ( | BioLegend, cat#135252, RRID: AB_2910292 |

Table S5: Fluorochrome-conjugated antibodies used in staining of non-immune cells for flow cytometry

| | |
|---|---|
| CD45-BV750 | BioLegend, cat# 103157, RRID: AB2734155 |
| CD140a-PE-CF594 | BD Biosciences, cat# 562775, RRID: AB_2737786 |
| CD140b-APC | BD Biosciences, cat# 136008, RRID: AB_2268091 |
| CD31-PerCP/Cyanine5.5 | BioLegend, cat# 160206, RRID: AB_2910328 |
| Podoplanin-PE | BD Biosciences, cat# 566390, RRID: AB_2739721 |
| MAdCAM-1-BV421 | BD Biosciences, cat# 742812, RRID: AB_2741064 |
| PD-L1-BV711 | BioLegend, cat# 124319, RRID: AB_2563619 |
| LTbR-PE/Cyanine7 | BioLegend, cat# 134410, RRID: AB_2728153 |
